# Mechanistic basis of lineage restriction

**DOI:** 10.1101/2024.08.07.606262

**Authors:** Bohou Wu, Jae Hyun Lee, Kara M. Foshay, Li Zhang, Croydon J. Fernandes, Boyang Gao, Xiaoyang Dou, Chris Z. Zhang, Guoping Fan, Becky X. Xiao, Bruce T. Lahn

**Author notes:** To whom correspondence should be addressed. Email: Bruce Lahn,; Bohou Wu.

## Abstract

Lineage restriction is the biological phenomenon where cells lose developmental potency during differentiation for all except their adopted lineages. Lineage restriction is foundational to multicellularity as it secures the functional identities of the myriad cell types in the body. As yet, the mechanisms of lineage restriction remain enigmatic. By developing Potency-Seq to systematically map transcriptional potency of genes across the genome, we uncovered a link between the restriction of developmental potency of cells and the occlusion of transcriptional potency of their genomes. We further showed that, strikingly, genes can undergo irreversible occlusion of their transcriptional potency simply by chromatinization into the nucleosomal form with unmodified histones. These findings led to a comprehensive mechanistic account of lineage restriction as driven by gene occlusion. Specifically, naive pluripotent stem cells at the onset of development possess the unique capacity to erase occlusion globally to reset full transcriptional potency of the genome, which in turn establishes full developmental potency of the cells. Such deocclusion capacity is abolished when cells progress from naive to primed pluripotency. From this stage onward, lineage-inappropriate genes can undergo nucleosome-mediated occlusion in an irreversible manner when their cognate transcription factors (TFs) and/or placeholder factors (PFs) no longer exist in cells to protect them. Consequently, the transcriptionally potent portion of the genome shrinks progressively and irreversibly during differentiation, thus permanently restricting the developmental potency of differentiating cells.

**ONE-SENTENCE SUMMARY:** Developmental potency of cells is linked to transcriptional potency of their genomes

## INTRODUCTION

The hallmark of multicellular life is the presence, within a single organism, of a wide range of different cell types bearing the same genome but disparate transcriptional profiles. This is achieved when a single zygote proliferates and progressively differentiates along a branchwork of lineages to eventually give rise to a multitude of specialized cell types. A defining feature of this process is lineage restriction (aka cell fate restriction), whereby differentiating cells progressively lose their developmental potency for all except their adopted lineages (Fig. 1A) (*1*). Lineage restriction is a prerequisite for multicellularity as it ensures that the myriad cell types in the body, once formed during development, would faithfully maintain their committed functional identities over the entire lifespan of the organism (*1, 2*). As yet, the mechanisms of lineage restriction are essentially unknown and remain one of the most important open questions in biology (*2, 3*).

**Figure 1.**
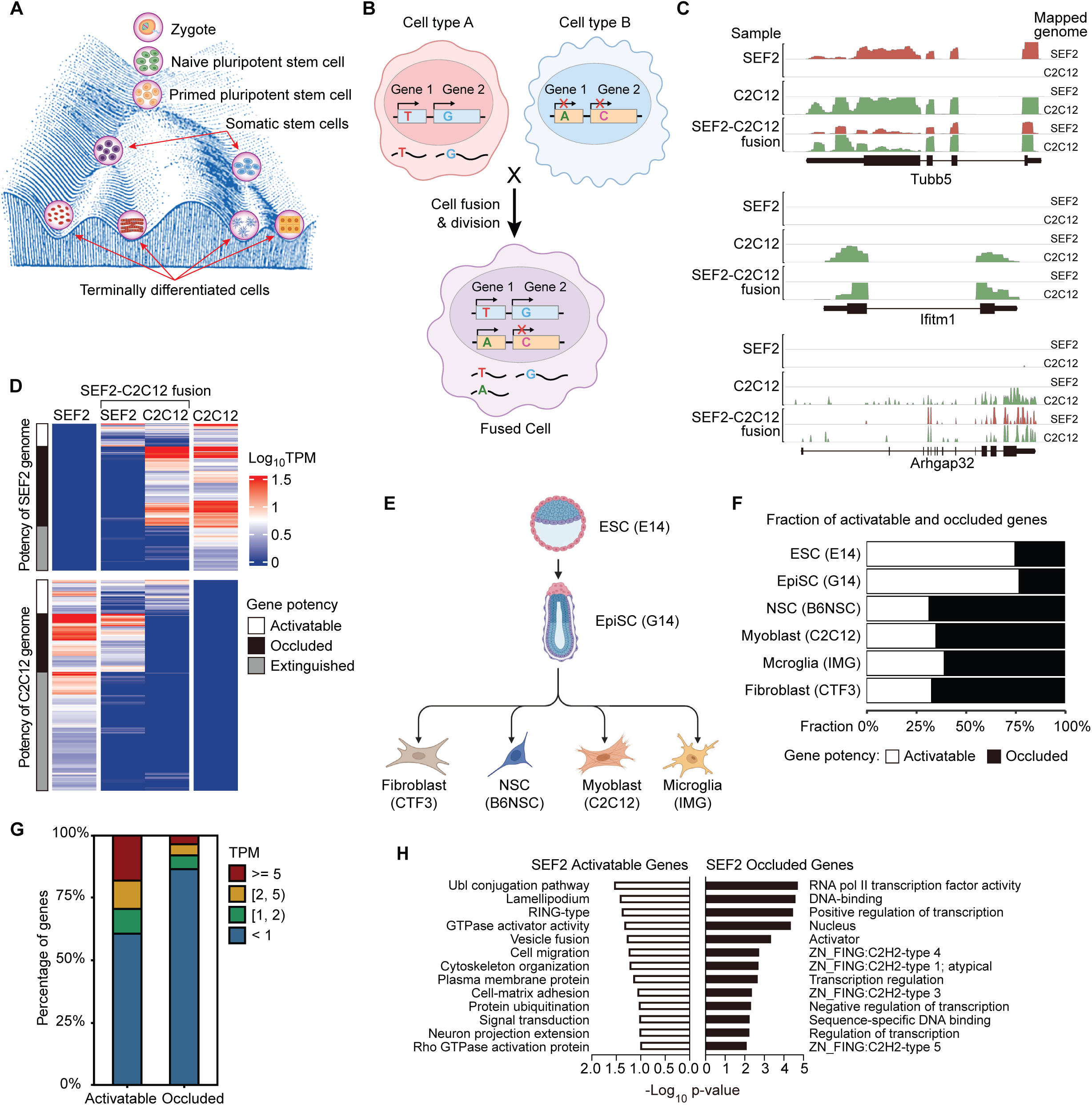
Restriction of developmental potency of cells is associated with reduction in transcriptional potency of their genomes. (**A**) Adaptation of Waddington’s classic illustration metaphorizing lineage restriction (ref. 1), with balls representing cells in different stages of development. It shows that differentiation is irreversible (the ball can only roll downhill), is accompanied by a progressive reduction of fate potency (possible paths remaining for the ball become progressively limited), and eventually reaches a stable differentiated state (the ball comes to a rest in one of the valleys at the bottom of the hill). (**B**) Schematic of Potency-Seq to measure gene potency in cell type B. In this cell, gene 1 possesses transcriptional potent as it is silent before fusion but activatable upon fusion, whereas gene 2 lacks transcriptional potency as it remains silent after fusion even though its ortholog in cell type A is actively expressed in the same fused cell. Sequence polymorphisms between the two cell types are indicated. (**C**) Potency-Seq profiles of representative constitutively expressed gene Tubb5, activatable gene Arhgap32, and occluded gene Ifitm1 in SEF2, before and after cell fusion. Only reads that can be mapped specifically to either the SEF2 or C2C12 genome based on inter-strain polymorphisms were used to build the profiles. (**D**) Gene expression heatmaps of unfused SEF2 (n = 303 genes) and C2C12 (n = 435), as well as fused cells. Shown are SEF2 or C2C12 genes classified as activatable, occluded or extinguished, based on their expression patterns pre and post-fusion. (**E**) Cell lines of varying developmental potency used in Potency-Seq. (**F**) Genome potency as reflected by the fraction of activatable and occluded genes in various cell lines. Genome potency of G14 (n = 427), NSC (n = 426), C2C12 (n = 190), and IMG (n = 652) was measured by fusion with SEF2. Genome potency of E14 (n = 362) and CTF3 (n = 110) was measured by fusion with, respectively, R1A (day 8 post-fusion) and C2C12. (**G**) Percentage of B6NSC occluded (n = 198) or activatable (n = 86) genes falling into high (TPM ≥ 5), medium (2 ≤ TPM < 5), low (1 ≤ TPM < 2) or no (TPM < 1) expression range in astrocytes differentiated from B6NSCs. (**H**) GO terms enriched in SEF2 occluded and activatable genes identified in SEF2-NSC fusion.

Cell fusion has been employed to investigate lineage-specific gene expression (*4–8*). Here, we further refined the cell fusion assay into a method termed Potency-Seq to measure the transcriptional potency of silent genes across the genome based on whether they can be activated by their cognate TFs. This assay revealed that genes indistinguishably silent in transcriptional activity can differ dramatically in transcriptional potency. We used “activatable” and “occluded” as binary measurements for the presence and absence, respectively, of transcriptional potency in silent genes. Activatable genes still possess the potency to be activated by their TFs, while occluded genes have lost such potency. We further demonstrated that, surprisingly, genes can become occluded by default simply by chromatinization into the nucleosomal form with unmodified histones, and that such nucleosome-mediated occlusion of transcriptional potency is irreversible even after DNA replication.

We employed Potency-Seq to systematically map genome-wide transcriptional potency of multiple cell types representing a wide range of developmental potency, and uncovered the following key mechanisms that link declining developmental potency of differentiating cells to reduced transcriptional potency of their genomes:

1. Naive pluripotent stem cells at the onset of development possess the ability to erase occlusion globally in order to restore full transcriptional potency across the genome, which in turn establishes full developmental potency of cells.
2. As naive cells progress into the primed pluripotent stage, such deocclusion capacity is abolished in preparation for differentiation, which is achieved via the downregulation of Esrrb, a key component of the deocclusion machinery. Notably, the silencing of Esrrb is itself conferred by occlusion. From this developmental stage onward, transcriptional potency of genes can only become unidirectionally occluded but not restored.
3. In primed pluripotent stem cells as well as somatic stem cells, PFs such as Sox2 protect silent genes whose activation is needed in later differentiation from nucleosome-mediated occlusion.
4. As differentiation proceeds down a specific lineage, genes whose expression is inappropriate to the adopted lineage, especially master regulators that drive alternative cell fates, undergo irreversible nucleosome-mediated occlusion at the opportune stage of differentiation when they lose protection from their TFs and/or PFs due to the disappearance of these factors from cells at such stage. This restricts the developmental potency of cells to within their adopted lineage because key genes that function outside these lineages are occluded and forever forgo the potency for activation.

Collectively, our results support a simple model of lineage restriction whereby the portion of the genome that remains transcriptionally potent shrinks progressively and irreversibly during differentiation, driving the developmental potency of differentiating cells to dwindle irreversibly as well. For convenience, we may refer to the developmental potency of cells just as cell potency, and transcriptional potency of genes or genomes just as gene potency or genome potency.

## RESULTS

### Assessing genome potency by Potency-Seq

Transcriptional potency of a silent gene can be assayed by whether it can be activated under various conditions such as stimulating the host cells with signal molecules, hypoxia condition, or induction of differentiation. However, the absence of gene activation may result from either the lack of gene potency or the unavailability of TFs supporting gene expression. To assess gene potency more conclusively, we devised the Potency-Seq method based on cell fusion and RNA-sequencing of fused cells (Fig. 1B). By fusing two disparate cell types, silent genes in one cell type are exposed to TFs from the other cell type in which their orthologs are actively expressed. If a silent gene turns on after fusion, then it is transcriptionally potent, as it is activatable by its cognate TFs provided by the fusion. In contrast, if a silent gene stays silent while its ortholog in the fusion partner’s genome is actively expressed, then it is occluded and lacks the potency to respond to TFs availed by the fusion. Importantly, Potency-Seq relies on fusing two cell types from the same species, followed by whole-transcriptome analysis that exploits sequence polymorphisms between the two cell types to assign transcripts of a given gene to the correct fusion partner. Compared to fusing cells from different species, this approach minimizes any interspecies incompatibility between TFs and their target genes.

We first established two mouse fibroblast clonal cell lines, SEF2 and CTF3, from SPRET/EiJ and CAST/EiJ strains, respectively. We then fused either of them to multiple cell lines from common laboratory mouse strains (i.e., C57BL/6, 129 or C3H) representing a wide range of developmental potency. SPRET/EiJ and CAST/EiJ are inbred wild mouse strains bearing sufficient sequence differences from common lab strains such that inter-strain polymorphisms can allow transcripts of most genes in hybrid cells to be assigned to the correct fusion partner.

As a proof of concept, we fused SEF2 with the mouse myoblast cell line C2C12 (strain C3H), and cultured the resulting SEF2-C2C12 hybrid cells for over two weeks to ensure that SEF2 and C2C12 genomes were combined into the same nucleus after multiple rounds of cell division, and that any mRNA remaining from the pre-fusion parental cells were mostly eliminated.

Potency-Seq was then performed on the cells to evaluate gene potency in both SEF2 and C2C12 genomes. It showed that silent genes in these cells, while indistinguishable in their lack of transcriptional activity, varied dramatically in their transcriptional potency. Activatable genes like Arhgap32 in SEF2, which was silent before fusion but became active after fusion just like its C2C12 ortholog, possessed the potency for transcriptional activation. Conversely, occluded genes like Ifitm1 in SEF2, which remained silent after fusion just like before fusion, even though its C2C12 ortholog was actively expressed in the same hybrid cells, lacked the potency to be activated by its TFs availed by the fusion (Fig. 1C). Potency-Seq on SEF2-C2C12 fusion thus allowed us to systematically classify the potency of silent genes in these cells into either activatable or occluded category (Fig. 1D). We note that cell fusion cannot inform on the potency of a silent genes if it is inactive in both cell types being fused. Additionally, a gene differentially expressed in the two parental cell lines before fusion may become silent in both genomes after fusion, a scenario known as extinction. Genes can become extinguished upon fusion due to reduced availability of relevant TFs or the action of repressors (*8, 9*). We will refer to silent genes in a cell type whose potency can be ascertained by the fusion assay as informative genes.

We next fused C2C12 with CTF3 to test whether potency measurement is consistent between different fusions (Fig. S1A). Of the 120 C2C12 occluded genes identified in the SEF2-C2C12 fusion, 29 were informative in the CTF3-C2C12 fusion, and of these, 28 were also found to be occluded. This result, together with additional experiments shown later, demonstrates that the annotation of occluded genes in a cell line is remarkably consistent when fused to different partner cell lines (except when fused to naive pluripotent stem cells as described below). That a minority of genes might appear occluded in one fusion but activatable in another could be due to several sources of uncertainty in the assay (see Supplementary Text).

We selected Myf5, a myogenic master regulator occluded in CTF3 but expressed in C2C12 for further validation. PCR products from gDNA or cDNA of CTF3-C2C12 hybrid cells were sequenced (Fig. S1B). While similar levels of Myf5 gDNA from SEF2 and C2C12 were detected based on polymorphism, Myf5 mRNA was only detected from C2C12 in the fusion sample, indicating that only the C2C12 copies but not the CTF3 copies of Myf5 were expressed in hybrid cells.

### Restriction of cell potency is associated with reduction in genome potency

We next selected multiple mouse cell lines representing a wide range of developmental potency and assayed transcriptional potency of their genomes by Potency-Seq (Fig. 1E). For pluripotent stem cells, we used the mouse embryonic stem cell (ESC) line E14 derived from the inner cell mass (ICM) of pre-implantation blastocyst and the mouse epiblast stem cell (EpiSC) line G14 derived from the epiblast of post-implantation embryo (Fig. 1E). ESCs are said to possess naive pluripotency representing an early stage of development not yet committed to differentiation, whereas EpiSCs are said to possess primed pluripotency representing a later stage of development already primed to differentiate (Fig. 1A) (*10*). RNA-Seq on E14 and G14 confirmed the expression of characteristic naive and primed markers, respectively. For somatic cells with more restricted fate potency, we chose the mouse neural stem cell (NSC) line B6NSC, microglia cell line IMG, and the above described C2C12 and CTF3. SEF2 was selected for fusion with most of these cell lines to assess their genome potency because of it has abundant sequence polymorphisms with them.

To examine the transcriptional potency of E14, G14, B6NSC, C1C12 and IMG genomes, informative silent genes in these cells were classified as occluded or activatable based on fusion with SEF2. Similar analysis was also performed on CTF3 based on its fusion with C2C12. For the four somatic cell types, B6NSC, C1C12, IMG, and CTF3, more silent genes were occluded than activatable (Fig. 1F, Fig. S2A). For the pluripotent G14, by contrast, far more silent genes were activatable than occluded (Fig. 1F, Fig. S2B). We performed the same analyses on clonal hybrid lines derived from the above bulk fusions, and obtained similar results (data not shown).

Measuring E14 genome potency in SEF2-E14 fusion was problematic because the vast majority of SEF2-specific genes (i.e., active in SEF2 but silent in E14) became extinguished in the SEF2 genome after fusion (Fig. S2C), presumably due to ESC’s strong reprogramming ability that robustly repressed SEF2-specific genes (see more description below). We reasoned that reprogramming is a gradual process and as such, E14 genome potency can perhaps be measured in early-stage fusion samples before reprogramming has taken full effect. We therefore reanalyzed time-course fusion data from our previous study where the rat fibroblast cell line R1A was fused with E14 (*7*). Unlike SEF2, R1A fusion efficiency with E14 was high enough to produce sufficient amounts of early-stage hybrid cells for RNA-Seq analysis. Similar to SEF2-E14 fusion, most R1A-specific genes became extinguished in R1A-E14 hybrid cells after long-term cell culture. However, large-scale gene extinction was not obvious for day 2, 4, and 8 post-fusion, enabling measurement of E14 genome potency in these early-stage fusion samples (Fig. S2D). Similar to G14 cells, most informative silent genes in the pluripotent E14 cells were activatable (Fig. 1F, Fig. S2D). Notably, our fusions might overestimate the fraction of occluded genes. In particular, occluded E14 genes annotated in R1A-E14 fusion may be due to regulatory incompatibility between rat and mouse. Thus, E14 genome potency was likely underestimated (see Supplementary Text).

It has been proposed that the fate potency of cells is actively acquired during development by pioneer transcription factors, rather than gradually restricted (*11–13*). To address this, we examined the expression of activatable and occluded genes in B6NSC after their differentiation into astrocytes (Fig. 1G). While 14% (12 out of 86) of B6NSC activatable genes are activated, only 2.5% (5 out of 198) of B6NSC occluded genes showed expression in differentiated astrocytes (p<0.0005, Fisher’s exact test). This observation argues that the loss of gene potency is essentially irreversible during somatic differentiation, and that the developmental potency of differentiating cells is irreversibly restricted rather than actively acquired. The small number of occluded B6NSC genes that turned on after differentiation could be due to uncertainty in the measurement of occlusion (see Supplementary Text). Additional evidence for the irreversibility of occlusion in somatic cells is described below.

### Occluded genes are enriched for developmental master regulators

During lineage differentiation, the activation of lineage-specific genes follows a hierarchical order, with master regulator genes (MRGs) that specify the target lineage – typically TFs – turning on first, which in turn activate downstream effector genes to produce physiological properties characteristic of the resulting cell type (*14*). We reasoned that if gene occlusion indeed serves the purpose of restricting the fate potency of a differentiated cell type, then it might act on MRGs more than effectors, as occlusion of MRGs should be sufficient to shut down entire lineage programs including their target effectors. Gene ontology (GO) analysis is in line with this prediction. Genes occluded in SEF2 identified by the SEF2-B6NSC fusion were highly enriched in MRGs of neural fates, including C2H2-type zinc finger proteins such as Zic1, Zic2 and Zic5, high mobility group (HMG) box domain proteins such as Sox1, Sox2 and Sox11, as well as homeodomain proteins such as Pou3f1 and Pou4f1. By contrast, activatable genes were more enriched for downstream effector functions, such as neuronal projection and extension, GTPase activation, and vesicle fusion (Fig. 1H). Similarly, occluded SEF2 genes identified in SEF2-C2C12 and SEF2-IMG fusions were enriched in transcriptional regulators and developmental signals (Fig. S3).

The above observation also sheds light on the phenomenon of transdifferentiation by ectopic expression of MRGs or cell fusion (*15, 16*). On the surface, such transdifferentiation seems to contradict the irreversibility of gene occlusion or cell fate restriction (*17*). However, there is no contradiction if the forced introduction of ectopically expressed MRGs by either transgenes or cell fusion are only turning on their target effectors that are activatable, which is sufficient to confer phenotypic similarities to alternative cell fates, while leaving occluded genes still occluded. Indeed, we repeated the classic experiment of converting fibroblasts into myotubes by ectopically expressing the myogenic master regulator Myf5 (*18*), and noticed that muscle-related genes activated in fibroblasts by the transgene were themselves activatable, whereas occluded genes, including endogenous Myf5, remained silent (*6*) (unpublished data).

### Capacity to erase occlusion is present in naive pluripotency but abolished in primed pluripotency preceding lineage differentiation

Early mammalian embryos possess pluripotent stem cells capable of differentiating into all somatic lineages and the germline. Their pluripotency is proposed to progress from naive to primed stage before embarking on lineage differentiation (*10, 19*). The biological significance of different pluripotency states remains unclear.

We hypothesized that the naive state (represented by E14) and the primed state (represented by G14) differ in how they regulate genome potency in order to serve their specific developmental needs. To test this, we performed hierarchical clustering to quantify changes in expression patterns before and after fusion, focusing on genes differentially expressed between partner cells before fusion. For somatic-somatic fusions (i.e., SEF2 fused to C2C12, IMG or B6NSC), the analysis showed that, strikingly, the correlation of expression patterns from the same genome before and after fusion was much greater than that between the two fusion partner genomes in the same hybrid cells (Fig. 2A). This observation is consistent with the role of genome potency occlusion in safeguarding cell identity. By contrast, in the SEF2-E14 fusion, expression of the SEF2 genome was reprogrammed to become quite similar to E14 (Fig. 2A).

**Figure 2.**
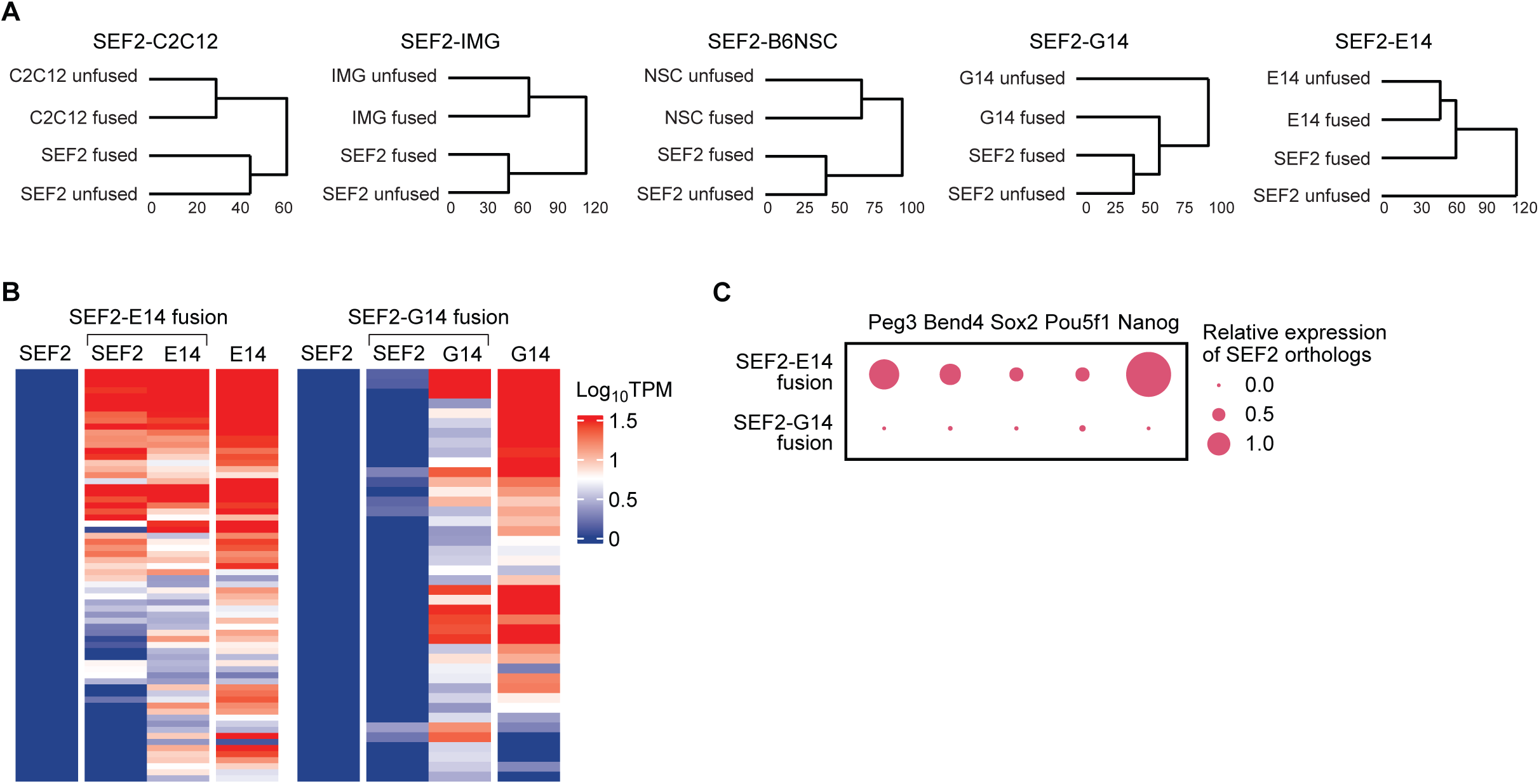
Deocclusion ability is present in naive pluripotency but lost in primed pluripotency. (**A**) Hierarchical clustering of pre-and post-fusion samples based on the expression patterns of genes differentially expressed between pre-fusion partner cells (TPM >= 2; log2 fold change > 2; SEF2-C2C12, n = 1308 genes; SEF2-IMG, n = 3164; SEF2-B6NSC, n = 2704; SEF2-G14, n = 3055; SEF2-E14, n = 3434). (**B**) Expression heatmap of SEF2 occluded genes with active orthologs in the pre-fusion partner cells. Depicted are SEF2-E14 (n = 68) and SEF2-G14 (n = 42) fusions (**C**) Relative expression of representative SEF2 occluded genes and pluripotency genes in SEF2-E14 and SEF2-G14 fusions. Peg3, Bend4 and Sox2 are identified as SEF2 occluded genes by SEF2-C2C12, SEF2-IMG and SEF2-B6NSC fusion, respectively. Expression levels from the SEF2 orthologs are scaled to that of their E14 or G14 fusion partner.

This is consistent with naive pluripotency’s reprogramming ability which, as described below, is due to its capacity to erase occlusion globally to reset full genome potency. Importantly, the SEF2-G14 fusion showed a very distinct pattern. Unlike E14, G14 did not reprogram the gene expression pattern of SEF2 after fusion, but rather adopted an expression pattern closer to SEF2 (Fig. 2A). This is consistent with G14’s high genome potency as described earlier (Fig. 1F), and low reprogramming ability as described below.

The above observation suggests that a crucial function of the transition from naive to primed pluripotency prior to lineage differentiation is to shut off the capacity to reset genome potency, such that any loss of genome potency in subsequent differentiation would be locked in place and not reversible. To test this hypothesis, we combined SEF2 occluded genes found in SEF2-C2C12, SEF2-IMG and SEF2-NSC fusions, and checked whether their potency can be restored in SEF2-G14 hybrid cells. Notably, these SEF2 occluded genes were rarely activated in any of the three somatic-somatic fusion pairs, confirming the stability of occlusion in somatic cells (Fig. S4). By contrast, upon fusion with E14, most SEF2 occluded genes were activated, indicating the remarkable deocclusion capacity of ESCs (Fig. 2B). Strikingly, when fused to G14, SEF2 occluded genes did not undergo massive activation. Instead, they mostly remained occluded similar to when SEF2 was fused to other somatic cells (Fig. 2B). This indicates that G14 does not possess the robust deocclusion capacity as E14.

We examined several representative SEF2 occluded genes that were actively expressed in both E14 and G14 genomes, including Peg3, Bend4, and Sox2. In the SEF2-E14 fusion but not the SEF2-G14 fusion, these SEF2 genes were robustly deoccluded, regaining expression levels comparable to their E14 orthologs (Fig. 2C). Similar findings were deduced for two pluripotency genes, Nanog and Pou5f1, whose potency in SEF2 was expected to be lost but was not ascertainable by fusion with other somatic cells. Nanog expression from the SEF2 genome fully resembled occluded genes, being activated upon fusion with E14 but not G14 (Fig. 2C). Pou5f1 was slightly activated from the SEF2 genome in SEF2-G14 hybrids, but to a much lesser extent as compared to the activation seen in the SEF2-E14 fusion (Fig. 2C). We performed the same analysis using clonal hybrid lines derived from the above bulk fusions, which produced similar results (data not shown). Collectively, the above data demonstrate convincingly that the ability to reset genome potency is present in pluripotent stem cells in the naive state, but shut off upon transition to the primed state before embarking on differentiation.

### Esrrb is responsible for the contrasting deocclusion ability between naive and primed pluripotency

While naive and primed pluripotent states share very similar gene expression patterns, there are a number of genes expressed in naive but silent in primed cells. We speculated that some of these genes, especially those implicated in pluripotency, might encode key components of the deocclusion machinery present in the naive state but absent in the primed state. We focused on two such candidate genes, Klf4 and Esrrb, both highly expressed in naive cells but silent in primed cells (*20, 21*), and which were previously implicated in promoting naive pluripotency (*22–26*). We fused either E14 or G14 with the rat neuroblastoma cell line B35 due to the ease to manipulate the resulting hybrid cells including lentiviral transduction. RNA-Seq on the G14-B35 fusion sample uncovered a set of robustly occluded genes in B35 in that they showed little expression from the B35 genome but high expression from the G14 genome of hybrid cells, and the same pattern persisted in two clones derived from the bulk fusion (Fig. 3A, left). In the E14-B35 fusion, by contrast, these occluded B35 genes showed noticeable activation by day 8 post-fusion, and even greater activation in two hybrid clones (Fig. 3A, middle). This is consistent with the presence of robust deocclusion capacity in naive but not primed pluripotent cells as described earlier.

**Fig 3.**
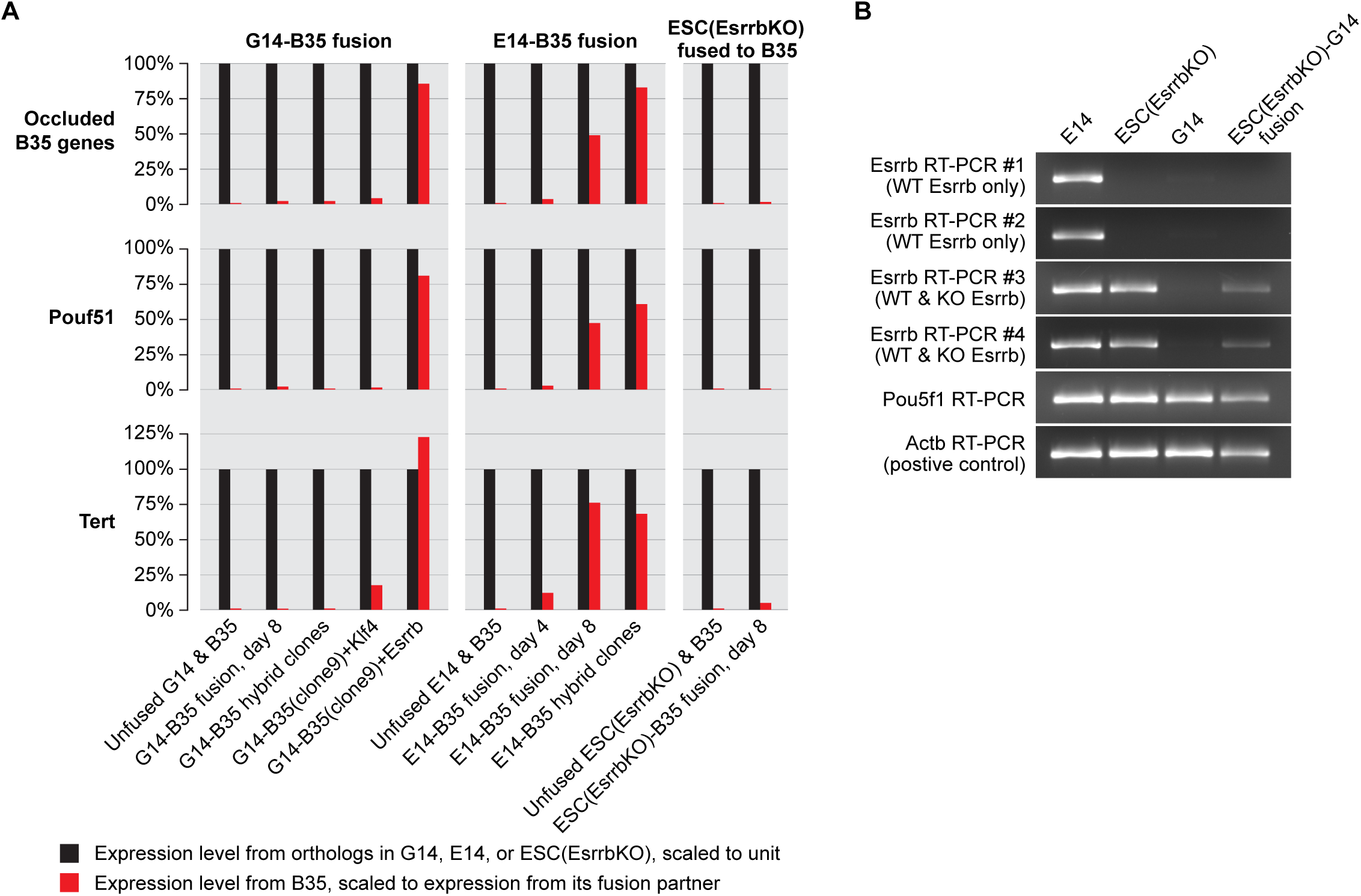
Deocclusion capacity is present in naive pluripotency, but abolished in primed pluripotency due to silencing of Esrrb by occlusion. (**A**) Expression levels of occluded B35 genes and their orthologs in fusions between B35 and various pluripotent cell types. (**B**) Cell fusion assay showing occluded status of Esrrb in G14. Esrrb expression was interrogated by four RT-PCR amplicons, two of which can only amplify the wildtype (WT) Esrrb transcripts, while the other two can amplify both WT and knockout (KO) Esrrb transcripts.

We then transduced the hybrid clone G14-B35(clone9) with lentivirus expressing either Klf4 or Esrrb, and performed RNA-Seq after prolonged culture. In the Klf4-transduced hybrid cells, the occluded B35 genes remained similarly differentially expressed between the B35 and G14 genomes, whereas in the Esrrb-transduced sample, the occluded B35 genes became robustly activated, reaching expression levels comparable to their G14 orthologs, just like in E14-B35 hybrid cells (Fig. 3A, left). Thus, expression of ectopic Esrrb but not Klf4 is sufficient to confer deocclusion capability to primed pluripotent cells that otherwise lack this ability.

Next, we examined whether Esrrb is necessary for the deocclusion capacity in naive cells. We utilized an Esrrb knockout ESC line in which exon 2 was deleted on both copies of the gene, referred to hereon as ESC(EsrrbKO) (*24*). RNA-Seq analysis confirmed that this cell line has a characteristic ESC transcriptome profile. ESC(EsrrbKO) was fused with B35, cultured for 8 days and harvested for RNA-Seq. For the occluded B35 genes, their behavior in this fusion was very similar to that of the G14-B35 fusion, namely they remained highly differentially expressed between the ESC(EsrrbKO) and B35 genomes of the hybrid cells just as in parental cells before fusion (Fig. 3A, right). This stands in sharp contrast to the fusion between B35 and the wildtype ESC line E14, demonstrating that Esrrb is necessary for the deocclusion capacity present in naive pluripotency. Collectively, the above data argue that the silencing of Esrrb in the primed state is responsible for the loss of the deocclusion capacity therein.

Two of the occluded B35 genes are noteworthy. One is the pluripotency gene Pou5f1, whose behavior is similar to that seen in SEF2 when fused to E14 or G14 as described earlier. The other is Tert, the telomerase gene known to be expressed in pluripotent stem cells and many cancers, but silent in most normal somatic cells (*27*). Our data suggest that Tert is not only silent, but occluded in somatic cells.

### Esrrb is occluded in primed pluripotent cells

We hypothesized that the silencing of Esrrb in primed pluripotent cells is itself conferred by occlusion. To test this, we fused ESC(EsrrbKO) to G14 to examine if the G14 copies of Esrrb would show signs of occlusion in fused cells. We used ESC(EsrrbKO) rather than a wildtype ESC line in the fusion for two reasons. First, any occluded genes in G14 would have their occlusion erased upon fusion with wildtype ESCs, given that the latter possess the deocclusion capacity. ESC(EsrrbKO) cells on the other hand, have lost their deocclusion ability due to Esrrb knockout as described above. Second, the Esrrb knockout allele in ESC(EsrrbKO) cells lacks exon 2 but retains other exons. It is therefore still expressed except its mRNA lacks exon 2 and is nonfunctional. This allows the mapping of Esrrb transcripts in hybrid cells to either the ESC(EsrrbKO) or the G14 genome based on whether it contains exon 2. We performed RT-PCR on the ESC(EsrrbKO)-G14 fusion sample after extended culture. Primers designed to only amplify the wildtype Esrrb transcripts did not produce any product, whereas primers designed to amplify both wildtype and knockout Esrrb transcripts produced strong products (Fig. 3B). This indicates that in hybrid cells, the wildtype Esrrb copies in the G14 genome is silent, whereas the knockout Esrrb copies in the ESC(EsrrbKO) genome is expressed, consistent with the occluded status of Esrrb in G14.

### Irreversibility of genome potency loss during lineage differentiation

We reasoned that in order for occlusion to play the role of restricting the developmental potency of somatic cells, it needs to act irreversibly even when cells encounter differentiation cues for alternative fates. To directly test this, we selected two clonal lines from each of the SEF2-E14 and SEF2-G14 fusions, and differentiated them into NSCs. Despite the existence of a fibroblast genome, all four hybrid clones successfully generated neurospheres (Fig. 4A). Less cell death occurred during the differentiation of the SEF2-E14 clones as compared to the SEF2-G14 clones, likely because the SEF2 genome in the SEF-E14 fusion but not the SEF-G14 fusion was extensively reprogrammed toward the pluripotent state as described earlier. In either case, typical neurospheres formed readily, which were plated as NSC monolayer and examined by RNA-Seq.

**Figure 4.**
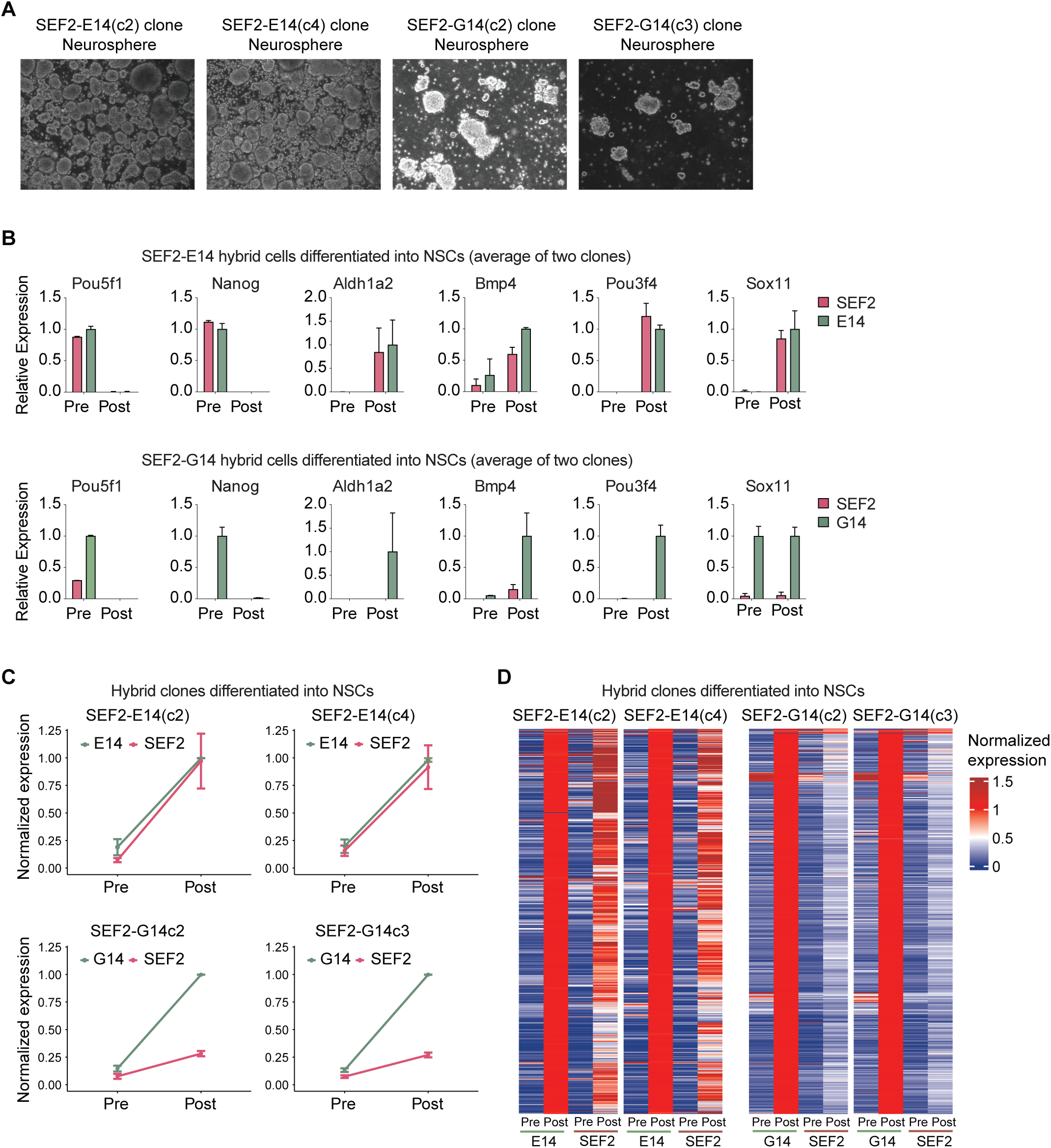
SEF2 occluded genes in SEF2-E14 but not SEF2-G14 hybrid cells possess the potency for proper activation in response to NSC differentiation. (**A**) Neurospheres successfully generated from the differentiation of four clonal hybrid lines, SEF2-E14(c2), SEF2-E14(c4), SEF2-G14(c2), and SEF2-G14(c3). (**B**) Expression of pluripotency genes and NSC markers before and after differentiation of SEF2-E14 and SEF2-G14 hybrid clones into NSCs (2 hybrid clones). (**C**) Average expression of occluded SEF2 genes whose orthologs were upregulated in the E14 or G14 genome of SEF2-E14 (n = 63 genes) and SEF2-G14 (n = 41) hybrid clones upon differentiation into NSCs. Expression level of each gene is scaled to the post-differentiation expression level of E14 or G14. (**D**) Expression heatmaps of upregulated genes (TPM >= 2, log2 fold change > 2, n = 889) in SEF2-E14 and SEF2-G14 hybrid clones following differentiation into NSCs. Expression level of each gene is scaled to the post-differentiation expression level of E14 or G14.

We first confirmed that the pluripotency markers Pou5f1 and Nanog turned off completely in all hybrid clones upon differentiation, indicating full exit from pluripotency (Fig. 4B). We then checked several NSC markers known to be upregulated when differentiating pluripotent cells into NSCs (Fig. 4B). In the two SEF2-E14 hybrid clones, these markers turned on from both SEF2 and E14 genomes after differentiation, consistent with the recovery of full genome potency of SEF2. Remarkably, in the two SEF2-G14 hybrid clones, whereas the NSC markers in the G14 genome were properly expressed in differentiated cells as would be expected, their SEF2 orthologs failed to turn on even as their G14 orthologs became properly expressed, indicating a permanent loss of transcriptional potency for these genes in SEF2. These results support the model that differentiating cells lose developmental potency via the irreversible occlusion of lineage-inappropriate genes, and are at odds with the notion that fate potency is actively acquired during development (*11–13*).

We next examined the overall expression of SEF2 occluded genes whose E14 or G14 orthologs were upregulated in the hybrids upon differentiation (Fig. 4C). As predicted by the unidirectional model of gene potency occlusion during lineage differentiation, expression of SEF2 occluded genes were only slightly elevated in SEF2-G14 hybrid clones post differentiation, in sharp contrast to their full activation in SEF2-E14 hybrid clones. To get a global view, we examined all the genes upregulated in the E14 genome of the SEF2-E14 fusion or the G14 genome of the SEF2-G14 fusion upon differentiation, irrespective of whether their potency was ascertained (Fig. 4D). In SEF2-E14 hybrid clones, many of these genes were upregulated to comparable levels from both SEF2 and E14 genomes upon differentiation into NSCs. By contrast, in SEF2-G14 hybrid clones, the great majority of genes showed full activation only in the G14 but not SEF2 genome. Essentially, albeit the two partner genomes in SEF2-G14 hybrid cells resided in the same nucleus, went through the same DNA replication, and experienced the same differentiation signals, only the G14 genome underwent proper differentiation. This result further contradicts the notion that cells actively acquire fate potency during development (*11–13*).

### In vitro assembled chromatin lacks transcriptional potency in somatic cells

We next explored the molecular basis of gene potency regulation. It is generally assumed that eukaryotic genes possess transcriptional potency by default, and their stable silencing requires additional repressive chromatin modifications (*28, 29*). Accordingly, we previously probed the mechanism of occlusion by searching for chromatin marks associated with occluded genes. But this approach yielded little success (*8, 30*), prompting us to consider an opposing model that genes could be occluded by default unless protected by some means from entering this default state. The eukaryotic nuclear genome is packaged by histones into nucleosome arrays as the default state, except in regions protected by DNA-binding factors such as TFs. DNA undergoing such packaging is said to be chromatinized. We therefore explored whether chromatinization alone was sufficient to render genes occluded by default.

Previous studies have shown that in vitro transcription is inhibited by chromatinization of the DNA template (*31, 32*). But whether chromatinized DNA possess transcriptional potency in living cells was never tested. To examine this, we selected four promoters, EF1A, Hsp68, UBC, and Nanog, and made four reporter plasmids each bearing one of the promoters driving a different fluorescence reporter (Fig. 5A). Plasmid DNA was chromatinized in vitro with HeLa core histones, or recombinant core histones lacking eukaryotic posttranslational modifications. Partial digestion of chromatinized DNA with MNase revealed regularly spaced nucleosome arrays, confirming successful chromatinization (Fig. 5B). We then transfected naked or chromatinized DNA constructs into 293T cells followed by three days of culture. Remarkably, while robust fluorescence was observed in cells transfected with naked constructs, only sparse and dim signals were detected in cells transfected with chromatinized constructs (Fig. S5). We quantified mRNA expression levels from the fluorescence reporters by RT-qPCR, and normalized it to the amount of plasmid DNA reaching the nuclei of transfected cells as measured by qPCR on DNA extracted from nuclei. Consistent with the fluorescence data, chromatinized constructs exhibited negligible expression compared to naked constructs (Fig. 5C). This result revealed that simply by chromatinization into the nucleosomal form, genes can become unresponsive to their cognate TFs. Crucially, the silencing effect of nucleosome formation holds true even when using recombinant histones devoid of any eukaryotic posttranslational modifications. This argues that nucleosomes can trigger gene silencing without any DNA or histone modifications, though it is reasonable to assume that modifications can subsequently be added to the silent chromatin (see further discussion below). We took these data as evidence that occlusion can be the default state of chromatinized DNA in somatic cells.

**Figure 5.**
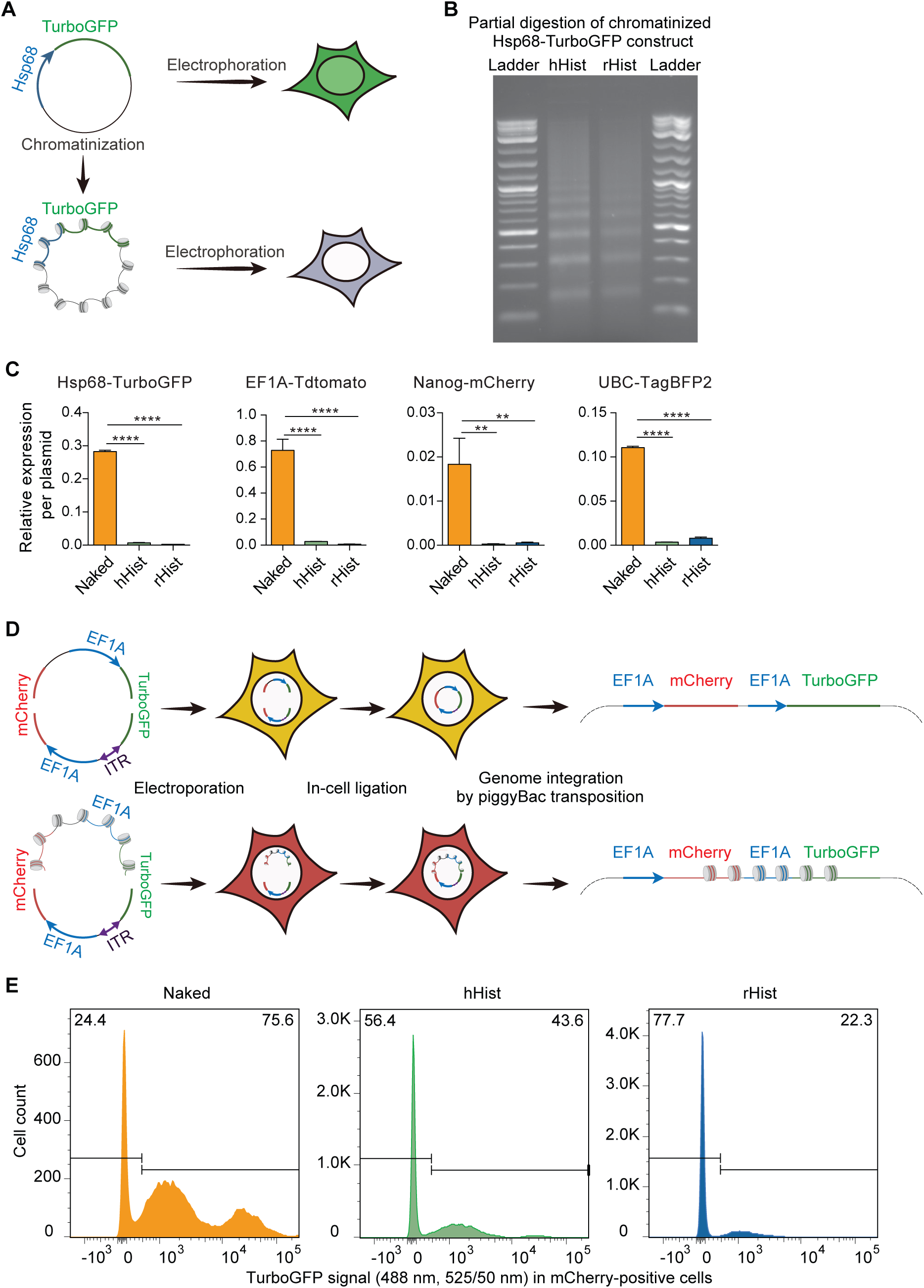
Chromatinized DNA undergoes occlusion by default, which is heritable through DNA replication. (**A**) Schematic of the chromatinization assay to measure transcriptional potency of naked or in vitro chromatinized DNA construct, using the Hsp68-TurboGFP construct as an example (PolyA downstream of reporter not shown). (**B**) Representative partial digestion by MNase of plasmid DNA construct chromatinized with HeLa core histones (hHist) or recombinant core histones (rHist). (**C**) Expression of naked and chromatinized constructs driven by different promoters in 293T cells. Reporter expression was measured by RT-qPCR and normalized to the amount of plasmid DNA extracted from cell nuclei as measured by qPCR, then scaled to endogenous Actb expression (3 replicates, ordinary one-way ANOVA test). (**D**) Schematic of the assay involving half-chromatinization followed by genome integration (PolyA downstream of reporter not shown). Expression cassettes of TurboGFP and mCherry were split within their coding sequences. The TurboGFP fragment in either naked or chromatinized form was co-electroporated into 293T cells with the naked mCherry fragment. The two fragments ligated spontaneously after entering cells to reconstitute a circular construct with complete TurboGFP and mCherry expression cassettes. The piggyBac ITRs in the naked mCherry fragment enabled the reconstituted construct to integrate into the host genome by piggyBac transposition. (**E**) Histograms of TurboGFP fluorescence in

### Nucleosome-mediated gene occlusion can occur at the resolution of individual genes

One caveat of the above experiment is that the expression difference observed between chromatinized and naked plasmids might be due to the former being more prone to getting trapped during electroporation in areas of the nucleus unconducive to transcription. Additionally, it does not speak to the resolution of nucleosome-mediated transcriptional repressive, namely, whether it impacts individual genes or expansive regions spanning multiple genes. To address these questions, we devised a half-chromatinization assay. We prepared two reciprocal linear DNA fragments with compatible sticky ends to facilitate their ligation into a circular construct (Fig. S6A). The sticky ends were designed to only permit ligation between the two fragments in the correct orientation, and did not support circularization of either fragment or ligation between the two fragments in the wrong orientation. One fragment carried the 3’ half of the mCherry reporter with a polyadenylation signal (PolyA), followed by the EF1A promoter driving the 5’ half of the TurboGFP reporter. It was termed the TurboGFP fragment because its promoter activity would be reflected by TurboGFP expression. The other fragment carried the 3’ half of TurboGFP with a PolyA, followed by the EF1A promoter driving the 5’ half of mCherry, and was termed the mCherry fragment. These two reciprocal DNA fragments complement each other in that, individually, neither could express any functional TurboGFP or mCherry due to their truncation, but when the two fragments were ligated together, the result was a circular reporter construct containing complete expression cassettes for both TurboGFP and mCherry. We chromatinized either the TurboGFP or mCherry fragment with HeLa histones, and ligated it to the naked version of the reciprocal fragment (Fig S6A). In theory, each ligated DNA molecule should be half-chromatinized and half naked. Partial digestion with MNase showed that the pattern of regularly spaced nucleosome array was still visible, but diluted by a background smear as would be expected from the presence of naked DNA and possibly also nucleosome sliding (Fig. S6B). To examine ligation fidelity, we performed PCR across the ligation junctions and sequenced the PCR products, which produced the correct sequences expected from proper ligation. We then transfected the ligated samples into 293T cells, and observed that the ligation product between chromatinized TurboGFP and naked mCherry gave rise to an enrichment of mCherry single-positive cells, whereas the opposite was true for the ligation product between naked TurboGFP and chromatinized mCherry (Fig. S6C). RT-qPCR was used to quantify mRNA expression levels from the TurboGFP and mCherry reporters, and results were normalized to the amount of ligated DNA reaching the nuclei of transfected cells as measured by qPCR on DNA extracted from isolated cell nuclei. PCR primers were designed to flank the ligation junctions in order to only interrogate successfully ligated DNA. Consistent with fluorescence data, the ligated construct bearing chromatinized TurboGFP fragment showed reduced expression than the construct carrying the naked TurboGFP fragment, and the same is true for the mCherry fragment (Fig. S6D). The expression difference between chromatinized and naked DNA seen in this assay is not as dramatic as that seen in Fig. 5C, possibly due to nucleosome sliding on the ligated construct (*33*), which could partially degrade the difference in nucleosome loading between the chromatinized half and the naked half of the circular DNA molecule. This notwithstanding, the result lends additional support to our hypothesis that nucleosome formation alone can lead to gene occlusion by default. It also argues that the silencing effect of nucleosomes can occur at the resolution of individual promoters.

### Nucleosome-mediated gene occlusion is heritable through DNA replication

So far, the observation that nucleosomes alone could trigger gene silencing by default was made with episomal DNA that cannot replicate. To address if DNA replication could alter this effect, we sought to integrate the half-chromatinized DNA into the host genome. In the course of our work, we noticed that the two reciprocal DNA fragments used in the half-chromatinization assay could be transfected into cells without prior in vitro ligation, and upon cell entry, they would correctly ligate with each other to form a complete reporter construct, likely due to DNA repair mechanisms in cells (Fig. S7). We therefore co-transfected into 293T cells the naked mCherry fragment that also carried a pair of piggyBAC inverted terminal repeats (ITRs) (Fig. 5D), plus the TurboGFP fragment in either naked form or chromatinized with HeLa or recombinant core histones, along with a plasmid expressing piggyBac transposase to drive genome integration of the construct formed by in-cell ligation of the two reciprocal fragments. Partial digestion by MNase confirmed proper nucleosome array formation of the chromatinized TurboGFP fragment used for transfection (Fig. S8A). A week after transfection, mCherry-positive cells, enriched for piggyBac-mediated integration of in-cell ligated dual-fluorescence construct, were isolated by cell sorting. Cells were then cultured for over a month with continuous proliferation and assayed for fluorescence reporter expression (Fig. S8B). At this point, cells that remained mCherry positive should carry stably integrated reporter construct in the genome. We observed that when the TurboGFP fragment used in transfection was in the naked form, the great majority of mCherry-positive cells were also positive for TurboGFP. By contrast, when the TurboGFP fragment was chromatinized, the majority of mCherry-positive cells were negative for TurboGFP, and the effect was particularly pronounced when recombinant histones devoid of any posttranslational modifications were used (Fig. 5E).

Furthermore, the expression patterns of the reporters persisted in culture over time. These results further support our hypothesis that nucleosome formation alone could render genes occluded by default in somatic cells, and that either the occluded or expressed state, once established, is stably inherited through cell division. Inferring from this, we propose that during somatic differentiation, genes can become occluded simply by the disappearance of their TFs (and PFs as described below) that leaves them no longer protected from the default silencing effect of nucleosomes.

We note that in the chromatinized TurboGFP samples, a minority of mCherry-positive cells were TurboGFP positive, which could be due to nucleosome sliding that partially eroded the chromatin difference between the TurboGFP and the mCherry halves of the ligated construct.

Interestingly, the opposite was also observed in the naked TurboGFP sample, namely, a minority of mCherry-positive cells were TurboGFP negative. This suggests that the naked reporter could occasionally undergo occlusion. We speculate that there is competition between the transcription machinery and the nucleosome assembly machinery. It has been reported that naked DNA are readily assembled into chromatin when transfected into cells (*34*). We suggest that after our naked reporter construct enters cells, the transcription machinery tends to win the competition as TFs can bind to their target promoter on the construct with faster kinetics than nucleosome assembly. But occasionally, nucleosome assembly occurs first before productive TF binding could take place, leading to occlusion of the reporter. However, when the reporter construct is already chromatinized before entering cells, even if incompletely due to technical reasons or erosion by nucleosome sliding, the balance would tip heavily against TF binding and toward nucleosome-mediated occlusion. Importantly, either the expressed state or occluded state, once established for a gene, is self-sustaining even through DNA replication.

### Sox2 and Olig2 act as placeholder factors maintaining transcriptional potency of target genes

Our data thus far support the model that during somatic differentiation, fate potency is progressively restricted through the irreversible occlusion of lineage-inappropriate genes. Our data also point to a possible mechanism of occlusion, namely, genes can, upon the disappearance of their cognate TFs, become occluded simply through the default silencing effect of nucleosome formation in regions previously occupied by the TFs. However, our model presents a dilemma for lineage-specific genes silent in early development but turned on later to drive the differentiation of specific lineages. If occlusion is irreversible in somatic cells as we postulated, then such genes must not undergo occlusion when silent during early development, such that they can undergo either activation or occlusion in later differentiation. How then do lineage-specific genes retain transcriptional potency while silent? To resolve this dilemma, we postulated that stem cells in and beyond the primed pluripotency stage, including both primed pluripotent stem cells and somatic stem cells, utilize PFs to protect silent genes needed for later activation from premature occlusion. PFs should be similar to TFs in that they can bind target genes to shield them from nucleosome-mediated default occlusion, but they differ from TFs in that they do not drive transcription themselves. Identifying such PFs would fill a critical missing piece in our model.

We reasoned that should PFs exist, they might confer a more open chromatin configuration around their binding sites by displacing nucleosomes in a manner similar to TFs (*35, 36*).

Consequently, activatable genes might show greater chromatin openness if they are indeed enriched for PF binding. We tested this in NSCs by analyzing published ATAC-Seq data (*37*), which, when combined with our own gene potency data, allowed the profiling of chromatin accessibility of NSC activatable and occluded gene (Fig. 6A). While both types of genes were similarly silent in NSCs, activatable genes indeed showed greater openness, suggesting the presence of PF binding. Actively expressed genes showed much greater openness as expected.

**Figure 6.**
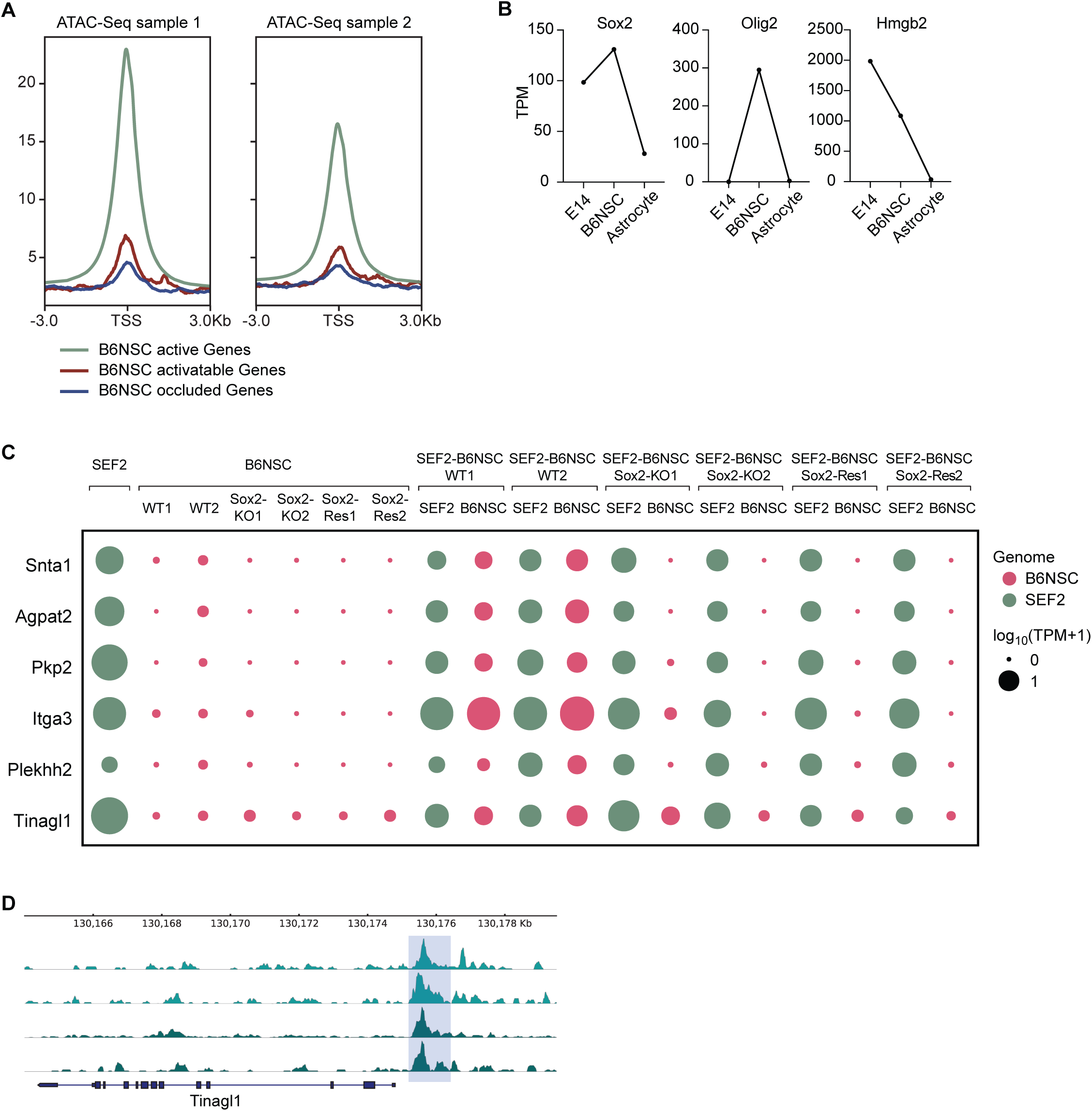
Sox2 acts as a PF to protect target genes from occlusion in NSCs. (**A**) ATAC-Seq profiles corresponding to activatable (n = 85) and occluded (n = 186) genes in B6NSC. (**B**) Expression of Sox2, Olig2 and Hmgb2 in E14, B6NSC, and B6NSC-differentiated astrocytes. (**C**) Bubble plot depicting expression levels in pre-and post-fusion SEF2 and B6NSC for activatable genes in wildtype B6NSC that became occluded following Sox2 knockout. Two B6NSC clones homozygous for Sox2 deletion were fused to SEF2 either directly (Sox2-KO1 and Sox2-KO2), or rescued for Sox2 expression by a lentivirus vector expression Sox2 before cell fusion (Sox2-Res1 and Sox2-Res2). Two wildtype B6NSC clones (WT1 and WT2) were used as control. (D) Sox2 binding profiles of a representative B6NSC activatable gene that became occluded following Sox2 knockout. Sox2 binding was measured by CUT&Tag with a monoclonal antibody (top two tracks for two replicates) and a polyclonal antibody (bottom two tracks for two replicates).

We reasoned that PFs should be abundantly expressed in NSCs where they bind to – and keep activatable – certain silent genes whose activation are required in later differentiation, but they should also turn off upon differentiation to allow their target genes to become either active or occluded. We therefore focused on three genes encoding DNA binding proteins previously implicated in neural development, Sox2, Olig2 and Hmgb2 (*38–42*), which are highly expressed in B6NSC but downregulated upon differentiation into astrocytes (Fig. 6B). We used CRISPR to delete coding regions of these genes in B6NSC, and fused the knockout cells with SEF2 to assay for changes in gene potency. Notably, in Sox2 knockout, six silent genes in B6NSC switched their potency from activatable to occluded status (Fig. 6C), indicating the requirement of Sox2 in maintaining the transcriptional potency of these silent genes. Importantly, reintroduction of Sox2 into knockout cells by lentivirus failed to revert the occluded status of these genes as measured by fusion with SEF2 (Fig. 6C), which is consistent with the irreversible nature of occlusion. These results argue that Sox2 act as a PF to protect these silent genes from becoming prematurely occluded in B6NSC. Moreover, the data suggest that genes undergoing occlusion after losing protection from their cognate PFs permanently lose responsiveness to such PFs, the same way that genes permanently lose responsiveness to their cognate TFs once occluded. Interestingly, three genes originally active in B6NSC cells became silent as well as occluded after Sox2 knockout (Fig. S9A), suggesting that Sox2 is required for maintaining not only their expression, but also their transcriptional potency.

To examine whether the Sox2 protein binds directly to these genes, we performed CUT&Tag (*43*), using several Sox2 antibodies. Of the above nine newly occluded genes following Sox2 knockout, reliable Sox2 binding peaks were detected in the vicinity of seven genes in wildtype B6NSC (Fig. 6D; Fig. S9D, Fig. S10).

We note that the real number of genes that lost transcriptional potency following Sox2 knockout should be much greater than what we observed. This is because ascertaining the transcriptional potency of silent genes in B6NSC, with or without Sox2 knockout, requires that the SEF2-B6NSC fusion is informative, namely, the genes are expressed from the SEF2 genome both before and after fusion. For genes silent in SEF2 to begin with, or active in SEF2 before fusion but become extinguished after fusion, the assay cannot speak to the potency of their B6NSC copies. The same applies to the other two NSC-related TFs analyzed below.

In Olig2 knockout B6NSC, we did not find reliable examples of activatable genes becoming occluded. However, 22 genes originally active in B6NSC lost both their expression and potency (Fig. S9B). We interpret this as Olig2 acting as a key TF to drive the expression of these genes, and once Olig2 was deleted, these genes not only became silent, but in the absence of Olig2 binding, underwent nucleosome-mediated default occlusion. There were also 10 genes originally active in B6NSC that turned off but remained activatable after Olig2 knockout.

Hmgb2-deleted B6NSC cells also did not show good examples of activatable-to-occluded switch. Of the seven genes originally expressed in B6NSC and turned off following Hmgb2 knockout, one became occluded while six remained activatable (Fig. S9C). Notably, Ier3 and Timp3 were silenced in both Sox2 and Hmgb2 knockout B6NSC. But their potency was lost only after Sox2 deletion (compare Fig. S9A with S9C). Consistent with this, Sox2 binding was detected in both genes (Fig. S9D), suggesting that while both Sox2 and Hmgb2 were required for the expression of these two genes, Sox2 likely served as a PF to maintain transcriptional potency of these genes even when they became silent upon Hmgb2 knockout.

Collectively, the above data support a placeholder model whereby primed pluripotent cells and somatic stem cells use PFs to sustain the transcriptional potency of silent genes that need to turn on in later differentiation. When these cells receive developmental cues to differentiate down a particular lineage, PFs disappear from cells and consequently, the genes they protect become either activated if TFs for these genes emerge, or occluded if otherwise. This model may underlie premarked or bivalent genes in stem cells, namely, silent genes thought to be poised for later activation that are bound by stemness factors, and/or possess both active and repressive histone modifications (*39, 44–46*). Mechanistically, PF binding may well be the causal agent of premarking and bivalency. Indeed, Sox2 binds to many inactive genes in NSCs that turn on during differentiation, and many Sox2-bound promoters are bivalent (*39, 47, 48*).

Importantly, the placeholder model is relevant to the long-explored but poorly understood concept of stemness (*49–52*) by arguing that whereas the source of stemness in naive pluripotent stem cells lies in their deocclusion capacity, the source of stemness in stem cells of later developmental stages lies in PFs that hold silent genes needed for subsequent activation in the activatable state.

## DISCUSSION

Our study provides a comprehensive mechanistic account of lineage restriction as follows. At the onset of development, naive pluripotent stem cells possess the deocclusion machinery that establishes full transcriptional potency of the genome, and in doing so, confers full developmental potency to cells. This puts cells on a blank slate, from which cells of more restricted transcriptional potency – and hence developmental potency – can be sculpted in subsequent differentiation. As naive pluripotent cells advance into primed pluripotency, said deocclusion capacity is abolished to prepare cells for lineage differentiation, and this is achieved via the occlusion of Esrrb, a key component of the deocclusion machinery. At this stage, the genome still retains full transcriptional potency, and cells retain full developmental potency, but genes can now undergo occlusion in ensuing differentiation. In primed pluripotent stem cells and also somatic stem cells, silent genes needed to turn on in later differentiation are protected from occlusion by PFs. As differentiation proceeds, developmental cues that drive cells to differentiate toward specific lineages would accomplish two things. One is turning on lineage-specific genes needed to specify target lineages; the other is occluding lineage-inappropriate genes no longer needed in the adopted lineages, especially MRGs for alternative fates. The latter comes about when cognate TFs or PFs of these genes disappear from cells at specific stages of differentiation. Upon losing protection from such factors, genes undergo irreversible occlusion via the default silencing effect of nucleosomes that have formed in regions previously protected by these factors. The overall outcome is a process that we previously hypothesized and termed “occlusis” (*5*), whereby the portion of the genome that retains transcriptional potency shrinks progressively and irreversibly during differentiation, pushing the fate potency of cells to dwindle irreversibly as well. Several outstanding questions about this process warrant further discussion.

One essential aspect of our model is that genes in somatic cells can undergo occlusion by the default silencing effect of nucleosomes when their cognate TFs or PFs become unavailable to protect them, and furthermore, once the occluded state is established, it is not reversible even when relevant TFs or PFs reappear in cells. How might DNA-histone and DNA-factor interactions facilitate such a behavior? We propose that genes can be bistable in that they can assume one of two energetically stable states (Fig. 7A). One is the factor-dominated state where genes are stably bound by trans-acting factors present in the cellular milieu (i.e., TFs or PFs). Genes in this state can be either active if the factors are TFs, or activatable if the factors are PFs. The other is the nucleosome-dominated state that corresponds to occlusion, where genes are stably packaged into the nucleosomal form. From a physicochemical perspective, these two states occupy two stable energy wells separated by a high energy barrier that prevents easy transition between the states (Fig. 7A). From an intuitive standpoint, DNA tightly wound in nucleosome arrays is shielded from factor binding, whereas DNA bound by an ensemble of factors on its cis-regulatory elements is shielded from nucleosome assembly.

**Figure 7.**
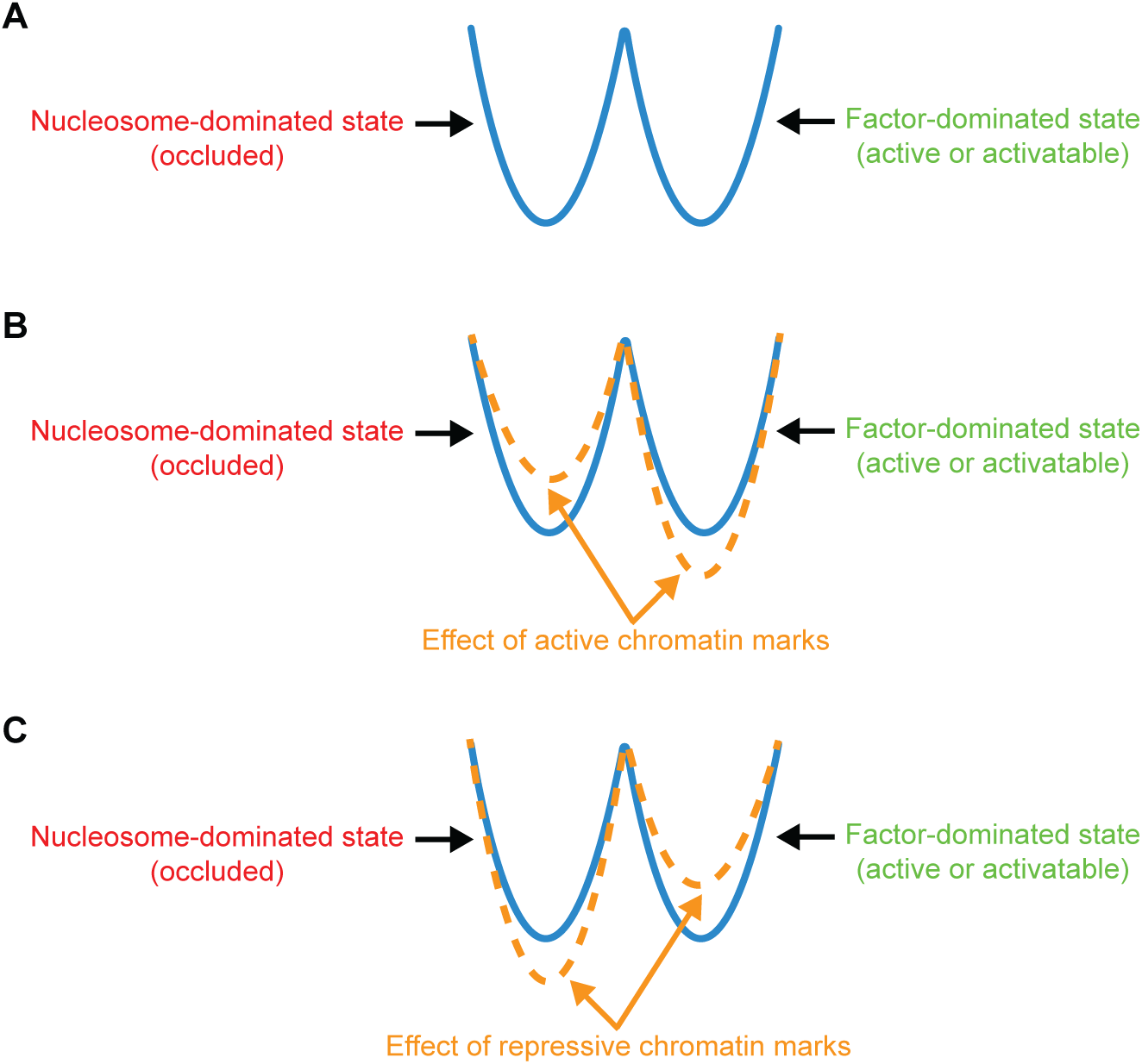
Bistability of genes and the modulating effect of chromatin marks on the bistability. (**A**) Genes can reside in one of two energetically stable states (represented by two energy wells) with a high energy barrier that prevents easy transition between them. In the nucleosome-dominate state, genes are occluded. In the factor-dominated state, genes are either active (if the factors are TFs) or activatable (if the factors are PFs). (**B**) Active chromatin marks increase the stability of factor-dominated state while decrease the stability of nucleosome-dominated state. (**C**) Repressive chromatin marks decrease the stability of factor-dominated state while increase the stability of nucleosome-dominated state.

Either of the two states, once formed, is self-sustaining and can perpetuate through DNA replication.

A large body of literature has implicated chromatin modifications in transcriptional regulation (*53*). Some modifications are associated with silent genes, such as DNA CpG methylation and histone H3K9 methylation, and are referred to as silent, or repressive, marks. Other modifications are associated with active genes, such as H3K4 methylation and H3K9 acetylation, and are referred to as active marks. What role might these marks play in gene occlusion? We argue that chromatin marks are not themselves the causal agent in establishing either the factor-dominated or nucleosome-dominated state of genes, as this task is accomplished by the appearance and disappearance of TFs and PFs during development as described above. Instead, chromatin marks serve the supporting role of reenforcing the stability of either state once established. For genes in the factor-dominated state, TFs (and potentially also PFs) can recruit active marks (i.e., H3K4 methylation and H3K9 acetylation) and remove silent marks (i.e., DNA methylation and H3K9 methylation), and in doing so, make DNA-factor binding energetically more favorable while DNA-histone interaction less favorable (*54, 55*) (Fig. 7B). Conversely, for genes in the nucleosome-dominated state, TFs are no longer present to recruit active marks, allowing silent marks to be added by a baseline machinery in the cell, which makes DNA-factor binding energetically less favorable while DNA-histone interaction more favorable (*56, 57*) (Fig. 7C). Thus, once either the factor-dominated or nucleosome-dominated state is established for a gene, that state can further induce the addition or removal of relevant chromatin marks to reenforce its stability, which in essence creates a deeper energy well that raises the barrier of transition to the opposite state. The end result of having these marks is reduced likelihood of genes undergoing spurious, unintended state transition. Indeed, we argue that the bistability between nucleosome-dominated state and factor-dominated state, and the role of chromatin marks in modulating this bistability, is the basis of many chromatin-related epigenetic phenomena. We also note that when chromatin marks are forcibly altered by artificial means, either globally or for specific genes (*8, 28*), the barrier between the two states are reduced, which could potentially allow some genes to undergo state transition.

Our study revealed that PFs such as Sox2 can keep silent genes in somatic cells activatable. It invites the question, do all activatable genes require PFs to stay activatable? We envision two scenarios. In the first one, some genes are “occludable”, meaning that they would undergo occlusion by nucleosome formation if not protected by their cognate TFs or PFs, but there are also genes that are “unoccludable”, meaning that they would not undergo occlusion even when chromatinized in the absence of factor binding. Unoccludable genes need not require PFs to stay activatable because their cognate TFs can overcome the energy barrier of nucleosomes to activate them. Unoccludable genes could include two types of genes that need not ever undergo occlusion. One is ubiquitously expressed housekeeping genes. The other is effector genes in terminally differentiated cells that are turned on by MRGs specific to those cell types. As discussed earlier, effectors don’t need to undergo occlusion because they will stay faithfully silent in lineages where they are not needed as long as their upstream MRGs are silenced by occlusion in those lineages. In the second scenario, all the genes in somatic cells require PFs to stay activatable even if they don’t ever need to undergo occlusion. In this case, housekeeping genes don’t require PFs because their TFs are present in all cell types at all times to drive their expression. But effector genes would need PFs to keep them activatable in stem cells whose later differentiation requires these genes to turn on. All considered, being unoccludable would seem to be a more parsimonious solution for effectors.

In summary, our study brings unprecedented mechanistic clarity to a fundamental and yet unanswered question in biology, namely, how does the fate potency of cells in multicellular organisms become irreversibly restricted during development. Additionally, the novel assays developed in this study could have broad future applications. Lastly, insights from the study could shed light on a number of burning questions beyond the context of lineage restriction, such as the mechanism of epigenetic silencing, the function of chromatin modifications, the source of bivalency of genes, the role of pioneer factors, and the essence of stemness. As such, our study has potential implications for many fields of biology, such as development, stem cell biology, gene regulation and epigenetics, and may also contribute to the understanding of disease processes such as developmental disorders, cancer and aging.

## MATERIALS AND METHODS

### Cell culture and fusion

Primary ear and tail fibroblasts were derived from adult SPRET/EiJ and CAST/EiJ mice, respectively, as previously described (*58*). Primary cells were passaged once and transduced with lentivirus expressing simian virus 40 large T antigen (SV40-T) to generate immortalized cells. They were then sorted into 96-well plates as single clones, giving rise to SEF2 and CTF3 clonal lines from SPRET/EiJ and CAST/EiJ, respectively. The mouse EpiSC cell line G14 (aka 1117E3) was derived from an E5.5 F1 embryo of a cross between C57BL/6 male and 129Jae female using published method and culture conditions (*20, 21*). Fluorescence and drug resistance markers were introduced into cells by lentivirus, after which another round of single-clone selection was performed to obtain the final clonal lines for cell fusion.

Cell lines and culture conditions were as described previously for C2C12 (from C3H mouse), B6NSC (C57BL/6 mouse) and, IMG (C57BL/6 mouse) (*59*), E14 (129 mouse) (*7*), ESC(EsrrbKO) (129 mouse) (*24*), and B35 (rat) (*8*). Briefly, SEF2, CTF3, C2C12, IMG and B35 were cultured in DMEM with 10% FBS. E14 and ESC(EsrrbKO) were cultured under feeder-free conditions in Knockout DMEM with 10% FBS, non-essential amino acids, sodium pyruvate, penicillin/streptomycin, β-mercaptoethanol, 3 uM CHIR99021 and 1 uM PD0325091. G14 was cultured on 10% FBS coated plates in DMEM/F12 supplemented with 20% Gibco Knockout Serum Replacement, GlutaMAX™, β-mercaptoethanol, penicillin/streptomycin, 12 ng/mL FGF2, 20 ng/mL ActivinA, 10 uM Y27632 and 2 uM IWP-2. B6NSC was cultured as monolayer in CELLstart substrate coated plates with DMEM/F12 supplemented with N2, B27, GlutaMAX, penicillin/streptomycin, 20 ng/mL FGF2 and 20 ng/mL EGF.

SEF2 or CTF3 were cultured in conditions of their fusion partners before fusion for a week as well as after fusion. The two cell lines to be fused were trypsinized, resuspended in medium, mixed thoroughly at 1:1 ratio, and plated into 6-well plates at high density to enhance cell-cell contact. Cells were settled for 2 hours to allow attachment to the plate, and treated with 45.5% PEG1000 pre-warmed to 42°C. After 1 minute of PEG treatment, cells were washed with fresh medium three times and cultured for 2 days. Hybrid cells were selected by dual drug selection or dual fluorescence-activated cell sorting. Chromosome loss could occur upon fusion, especially when clonal lines are derived from bulk hybrid cells (*8*), and genes on lost chromosomes could be wrongly annotated as occluded. To address this, we quantified strain-specific RNA-Seq reads for each chromosome in hybrid cells, and excluded fusion samples with signs of chromosome loss.

### In vitro differentiation

To induce astrocyte differentiation, B6NSC was seeded on poly-D-lysine coated 10 cm dishes at 5×10^5^ cells per plate, and cultured for 7 days with astrocyte differentiation medium containing DMEM/F12, N2, B27, penicillin/streptomycin, GlutaMAX™, 1 ng/mL EGF and 20 ng/mL BMP.

To induce NSC differentiation, SEF2-E14 or SEF2-G14 fusion cells were cultured in E14 or G14 conditions, respectively. One day before differentiation to NSCs, hybrid cells were passaged to allow 60%-70% confluency the next day. Upon differentiation, cells were washed with PBS to completely eliminate factors supporting ESC or EpiSC growth. Subsequently, hybrid cells were plated on 10 cm dishes coated with CELLstart at 1×10^6^ cells per plate, and cultured in N2B27 media containing DMEM/F12, Neurobasal Medium, N2, B27, GlutaMax, β-mercaptoethanol and 2 uM SB431542 to induce the neural fate. SEF2-E14 and SEF2-G14 fusion cells lost typical pluripotency morphology and acquired NSC morphology at day 5 and day 3, respectively. Cells were then dissociated by TrypLE Express Enzyme, and cultured in low attachment plates with NSC media containing FGF2 and EGF. Successfully differentiated cells would form neurospheres, which was purified by gentle centrifugation. Purified neurospheres were trypsinized and cultured as monolayers in CELLstart coated plates with NSC medium.

### RNA-Seq and strain-specific data analysis

Total RNA was extracted by MagNA Pure Compact RNA Isolation kit. Following DNase treatment, mRNA with polyA tail was purified with NEBNext poly(A) mRNA Magnetic Isolation module. Purified mRNA was reverse transcribed and the resulting cDNA made into libraries using Illumina primer sets following vendor’s protocol.

Over 30 million high-quality 2×150bp paired-end reads for each sample were obtained. The reads were aligned to N-masked mm10 mouse genome where SNPs between fusion partner genomes were replaced with the ambiguity base ‘N’. SNPsplit was used to extract reads specific to each strain. Transcripts per million (TPM) for each fusion partner was calculated based on the relative amounts of strain-specific reads. Genes in a given fusion partner were defined by pre-and post-fusion expression levels as activatable (pre-fusion TPM < 1, prefusion partner TPM ≥ 2, post-fusion TPM ≥ 30% of total post-fusion TPM of both genomes, average post-fusion TPM per genome ≥ 2), occluded (pre-fusion TPM < 1, prefusion partner TPM ≥ 2, post-fusion TPM < 10% of total post-fusion TPM of both genomes, average post-fusion TPM per genome ≥ 2), or extinguished (pre-fusion TPM < 1, prefusion partner TPM ≥ 2, average post-fusion TPM per genome < 2).

### Metagene analysis of SEF2 occluded genes in hybrid cells during differentiation

Strain-specific gene expression was calculated for SEF2-E14 or SEF2-G14 fusion cells before and after differentiation. Total TPM, calculated from all reads from both partners of hybrid cells, were used to select for activated genes (total TPM post-differentiation >= 4, fold change > 4) during NSC differentiation. For each gene, expression from the E14 or G14 genome post differentiation was scaled to unit. Expression from the E14 or G14 genome pre-differentiation, as well as expression from the SEF2 genome pre-and post-differentiation, was scaled to this unit.

### Chromatin assembly

Chromatinization was performed by Chromatin Assembly Kit from Active Motif (Cat #: 53500). The HeLa core histones in the kit were replaced with equal amounts of recombinant histones from NEB (Cat#: M2508S and M2509S) for recombinant histone samples. As a modification of the manual that improved chromatin assembly efficiency, high salt buffer and low salt buffer were mixed together during the incubation step of h-NAP-1 and core histones. Following chromatin reconstitution, 5 mM MgCl_2_ was added to the solution and centrifuged for 15 minutes to precipitate the assembled chromatin. This removed unchromatinized or partially chromatinized DNA. The pellet was resuspended with Gene Pulser electroporation buffer with 1 mM EDTA. Undissolved chromatin pellet was removed by centrifugation before electroporation.

### Partial digestion assay

Following purification of assembled chromatin, 3 ul of 0.1 M CaCl_2_ was added to 100 ul of resuspended chromatin. Subsequently, 1000 units of MNase was added to the solution and incubated at room temperature for 30 seconds. The digestion was stopped by adding 34 ul 4X Enzymatic Stop Solution in the Chromatin Assembly Kit. Following purification of digested DNA by QIAquick PCR Purification Kit from Qiagen, gel electrophoresis was used to visualize the pattern of nucleosome array.

### Cell transfection with naked or chromatinized DNA and measurement of gene expression

293T cells were transfected by electroporation. Cells were trypsinized and counted. One million cells were resuspended in Gene Pulser Electroporation Buffer containing 1 mM EDTA and 1 ug naked or chromatinized plasmid DNA (see below for plasmid ID used). Electroporation was conducted with Gene Pulser II Electroporation System using recommended parameters. MNase was added to the medium one-day post electroporation to remove naked or chromatinized DNA molecules that did not enter the cells. Reporter expression was visualized by fluorescence microscopy. To assay for promoter strength, electroporated cells were separated into two batches, one to extract mRNA for RT-qPCR quantification of reporter expression, and the other to isolate nuclear plasmid DNA for qPCR quantification. The mRNA quantity was normalized to the plasmid DNA quantity to obtain the final, normalized reporter expression levels. Chemical transfection (i.e., by lipofectamine) was not used due to the possibility that the chemicals used could disrupt DNA-histone association (*60*).

### CRISPR-mediated gene knockout

Guide RNAs were designed at 5’ and 3’ UTRs of Sox2, Olig2 and Hmgb2 genes to delete their full coding sequences. CRISPR plasmids were then constructed, each expressing an sgRNA pair corresponding the 5’ and 3’ UTR targets of a gene, plus Cas9 and blasticidin drug resistance (see below for plasmid ID used). Following transfection of plasmid DNA, B6NSC was selected by 30 ug/ml blasticidin for 2 days to enrich for cells that took in functional copies of the plasmid, and then sorted into CELLstart coated 96-well plates to derive clonal lines. The resulting clones were genotyped with primers across the two target sites to screen for successful deletion, and primers within the deleted fragment to screen for homogenous knockout. Knockout clones were further validated by RNA-Seq data, wherein the coding sequences of target genes were devoid of reads.

### Rescue of Sox2 knockout

A lentiviral vector containing the PGK promoter driving Sox2 as well as a neomycin resistance gene were created and packaged into virus by VectorBuilder (see below for plasmid ID used). Sox2 knockout B6NSC clones were transduced at MOI of 5 and selected by G418 for 7 days. The rescue of Sox2 expression was confirmed by RNA-seq data, wherein the read coverage of Sox2 coding sequence was recovered.

### Plasmids and lentiviral vectors

Plasmids/vectors for chromatinization assay: VB171220-1258gxn, VB200911-1183yfq, VB210716-1111vfu and VB210719-1129pad; half-chromatinization assay: VB220428-1064cuh, VB221020-1032pzd, and VB220428-1062bsm; CRISPR-mediated knockout of Sox2, Olig2 and Hmgb2, respectively: VB220823-1355ufr, VB220823-1356hth and VB220823-1358fhs; lentiviral labelling of cells: VB150915-10026 (EGFP and puromycin resistance); VB150925-10020 (mCherry and hygromycin resistance); lentiviral expression of SV40-T, Esrrb, Klf4 and Sox2, respectively: VB171106-1316rqy, VB180510-1202zrv, VB181219-1169xkg, VB230911-1130rye.

All plasmids were constructed and lentiviruses packaged by VectorBuilder, and their maps and sequences can be retrieved by the above vector IDs at https://vectorbuilder.com by following menu link “Design Vector”, then “Retrieve Vector Information”. For the half-chromatinization assay, VB220428-1064cuh or VB221020-1032pzd (with piggyBac ITRs) and VB220428-1062bsm were digested by BbsI to create the mCherry fragment and the Turbo-GFP fragment with compatible sticky ends, which was subsequently gel-purified for the chromatin assembly assay.

### CUT&Tag

CUT&Tag was performed with NovoNGS CUT&Tag 3.0 High-Sensitivity Kit for Illumina from Novoprotein following vendor’s instructions. Both monoclonal antibody from Cell Signaling (Cat #: 23064) and polyclonal antibody from Abcam (Cat #: ab97959) against Sox2 were used.

## ACKNOWLEDGMENTS

We thank Marcelo Nobrega, Heng-Chi Lee, Alex Ruthenburg and Guohong Li for scientific advice, Marcelo Nobrega for administrative support, and Austin Smith and Graziano Martello for providing the Esrrb knockout ES cells. Data on Esrrb were also posted in a previous bioRxiv preprint (DOI: 10.1101/2021.05.04.442547) not yet published in a peer-reviewed journal. This work was funded by VectorBuilder, Frontier Explorer Foundation, Department of Human Genetics at the University of Chicago, and Chicago Biomedical Consortium with support from Searle Funds at The Chicago Community Trust.

## AUTHOR CONTRIBUTIONS

Supervision, formulation of direction and acquisition of funding: BTL; Conceptualization and methodology: BW, JHL, KMF, BTL; Investigation, data analysis and visualization: BW, JHL, KMF, LZ, CJF, BG, XD, BXX, CZZ, GF, BTL.

## DATA AND MATERIALS AVAILABILITY

All data and materials are deposited or available upon request.

## SUPPLEMENTARY TEXT

Several sources of uncertainty exist in the assessment of transcriptional potency of genes by Potency-Seq. First, while occlusion is measured at the gene level, it might actually occur at the level of individual cis-regulatory elements (e.g., enhancers). Specifically, a gene could possess multiple enhancers, with only a subset being occluded in a given cell line while the others remaining activatable. Such a gene would appear occluded in fusions that only brought in TFs cognate to its occluded enhancers, but the same gene would appear activatable in fusions that brought in TFs cognate to its activatable enhancers. It is even possible that the same enhancer could be occluded in regards to one set of TFs but activatable in regards to another set of TFs. Second, the statistical power in measuring expression levels of genes could be limited if they are lowly expressed or have few polymorphic sites to facilitate strain-specific mapping of RNA-Seq reads from hybrid cells. This, together with the inaccuracy of RNA-Seq, could lead to stochasticity in determining the occluded or activatable status of genes, which could occasionally result in the same gene being annotated as occluded in one fusion but activable in another. Third, while we attempted to mitigate interspecies incompatibility by fusing cells of the same species, we did deliberately choose to fuse cells from relatively divergent mouse strains in order to maximally exploit inter-strain sequence polymorphisms to specifically map gene expression in hybrid cells to the two fusion partners. Some level of inter-strain incompatibility might still exist, similar to that observed for cis-acting expression quantitative trait loci (cis-eQTL) (*1*). As a result, some activatable genes could be misclassified as occluded if their TFs in hybrid cells failed to activate them to sufficient levels due to inter-strain incompatibly. Fourth, while our analysis excluded fusion samples showing signs of chromosome loss, it remains possible, though perhaps not common, that local mutations such as circumscribed deletions could knockout the expression of a gene, resulting in it appearing occluded. The latter two of the above cofounding factors would lead to the overestimation of occluded genes and the corresponding underestimation of activatable genes. For fusions between mouse and rat cells, the greater interspecies incompatibility would further exaggerate the overestimation of occluded genes. Furthermore, in early-stage R1A-E14 fusion samples, additional overestimation of occluded E14 genes was possible if there were remnant mRNAs from extinguished R1A-specific genes that made the corresponding E14 orthologs seemingly occluded. These caveats notwithstanding, the internal consistency of our data across different fusion experiments involving multiple cell types indicate a relatively high degree of reliability of the fusion assay in ascertaining occluded and activatable genes.

---

**Figure S1.**
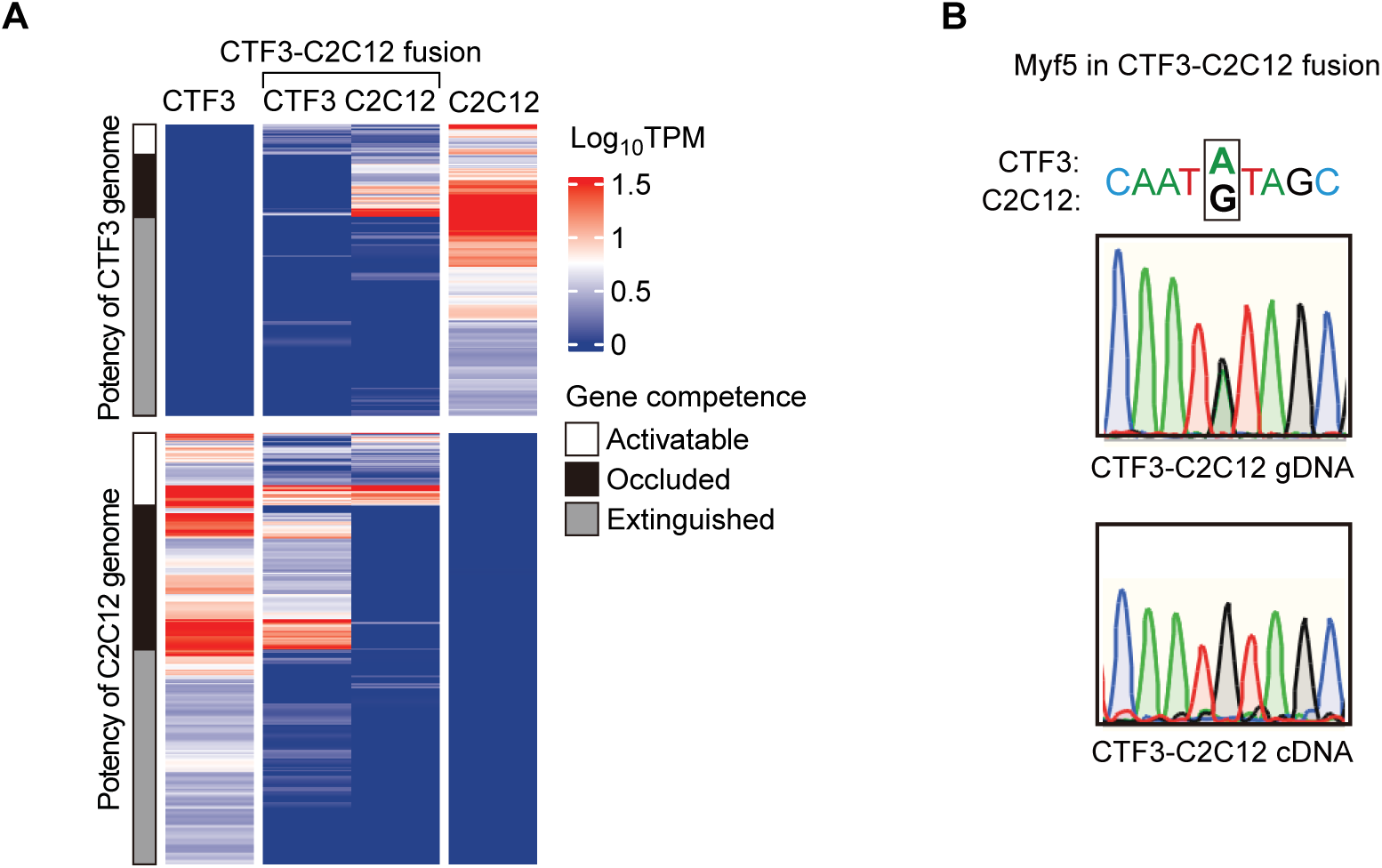
Potency-Seq of fused cells provides a measurement of gene potency. (**A**) Heatmap of informative silent gene (n = 858) expression in unfused CTF3 and C2C12 cells, as well as their strain-specific expression in CTF3-C2C12 hybrid cells. Activatable, occluded and extinguished genes are annotated based on their expression before and after cell fusion. (**B**) Sanger sequencing of RT-PCR (on CTF3-C2C12 cDNA) and genomic DNA PCR (on CTF3-C2C12 gDNA) products of Myf5, an occluded gene in CTF3, in CTF3-C2C12 hybrid cells. Signal at the SNP site reflects the relative abundance of Myf5 genomic DNA (showing two peaks on gDNA template) and mRNA (only one peak on cDNA template) in hybrid cells.

**Figure S2.**
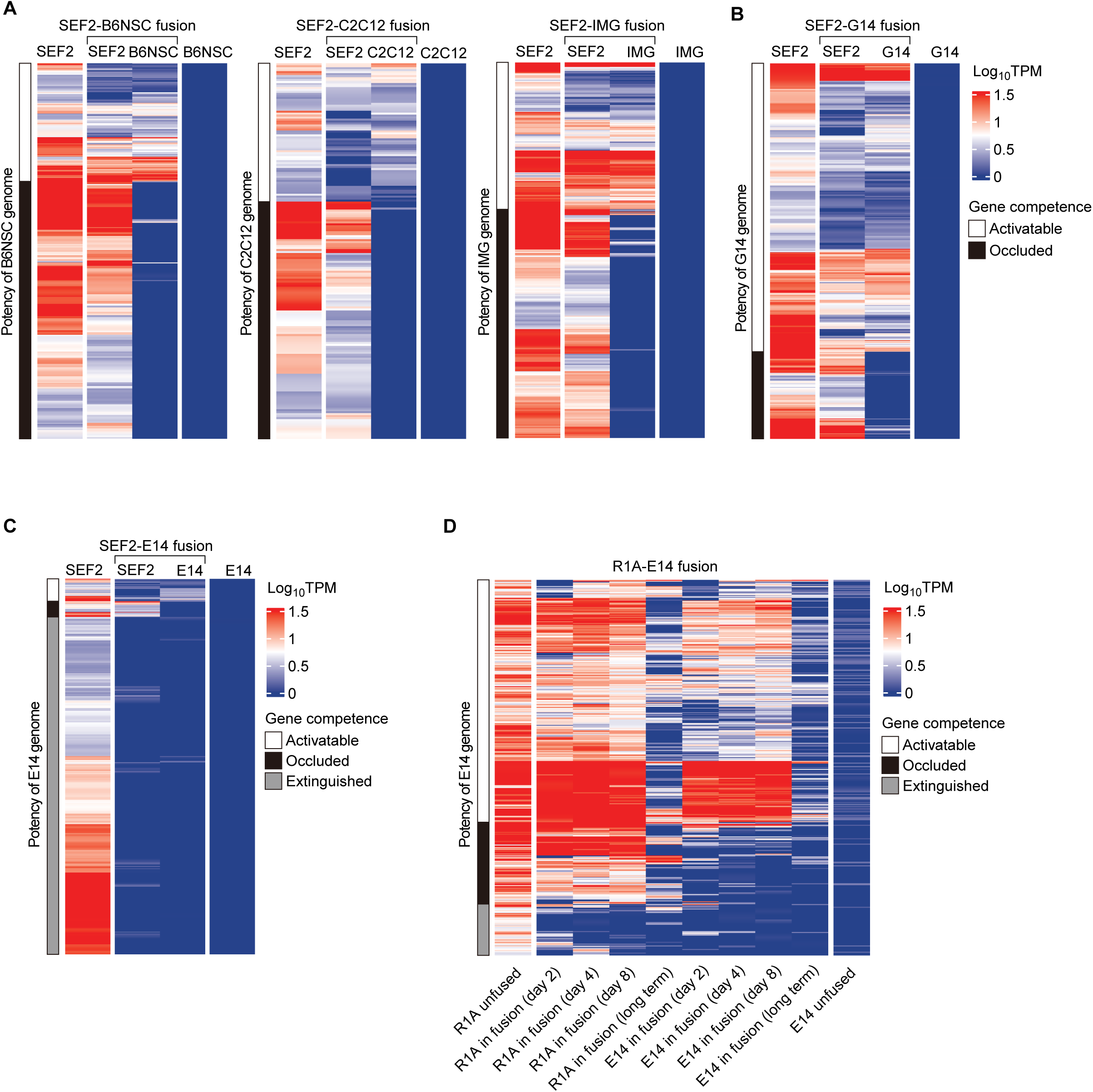
Genome potency of ESCs can be measured during early but not late stages of cell fusion. (**A**, **B**) Heatmap of informative silent gene expression in NSC (n = 426), C2C12 (n = 190), IMG (n = 652) and G14 (n = 427) unfused parental cells and their fusion cells with SEF2. (**C**) Heatmap of E14 informative silent gene (n = 788) expression in unfused SEF2 and E14 parental cells, as well as strain-specific expression in SEF2-E14 hybrid cells. (**D**) Time-course expression of E14 informative silent genes (n = 417) in R1A, E14 and R1A-E14 hybrid cells after cell fusion. Activatable, occluded and extinguished genes are annotated according to their expression before fusion and at day 8 post fusion.

**Figure S3.**
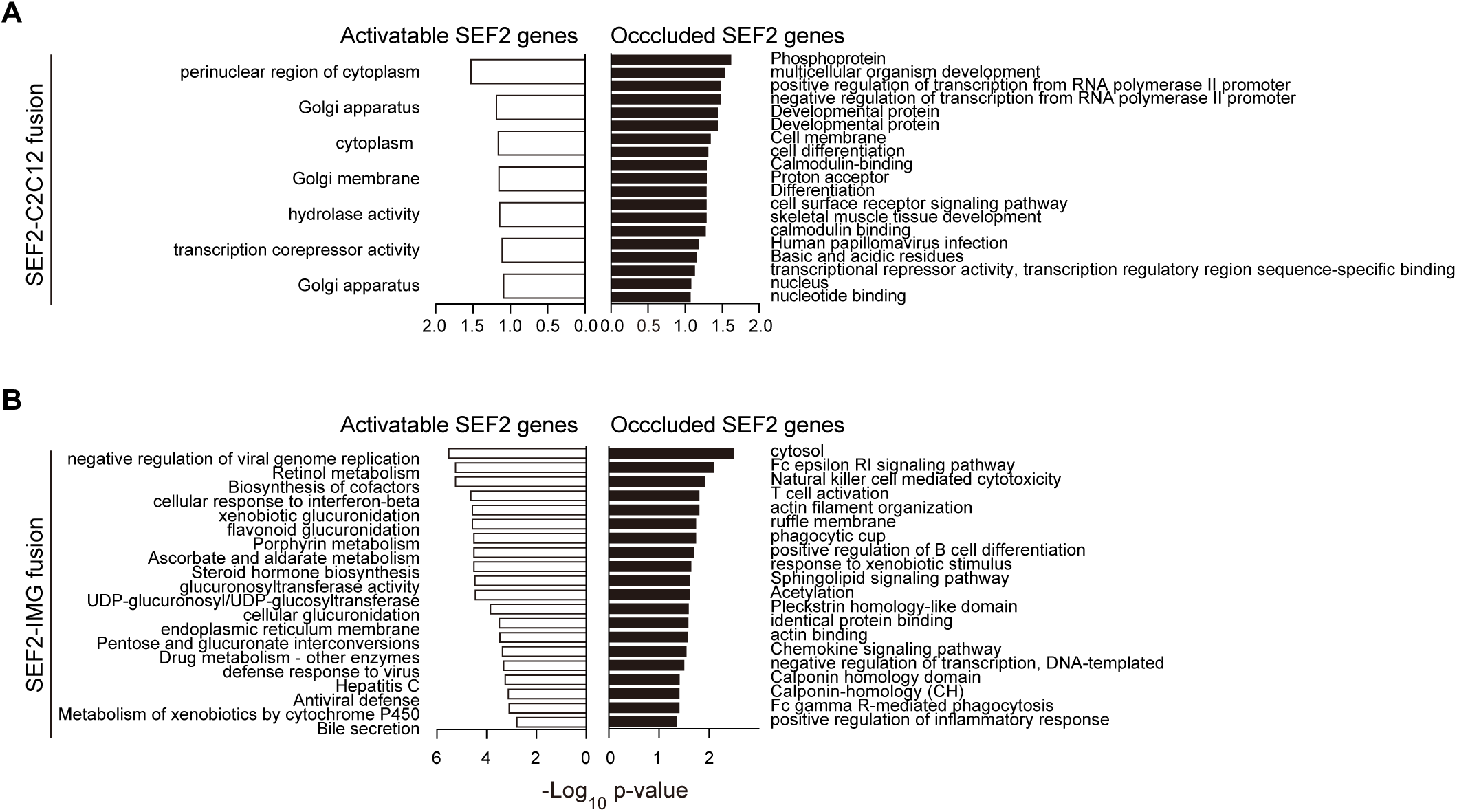
SEF2 occluded genes are enriched for regulatory functions. GO terms enriched in SEF2 activatable and occluded genes identified in SEF2-C2C12 (A) and SEF2-IMG (B) fusions.

**Figure S4.**
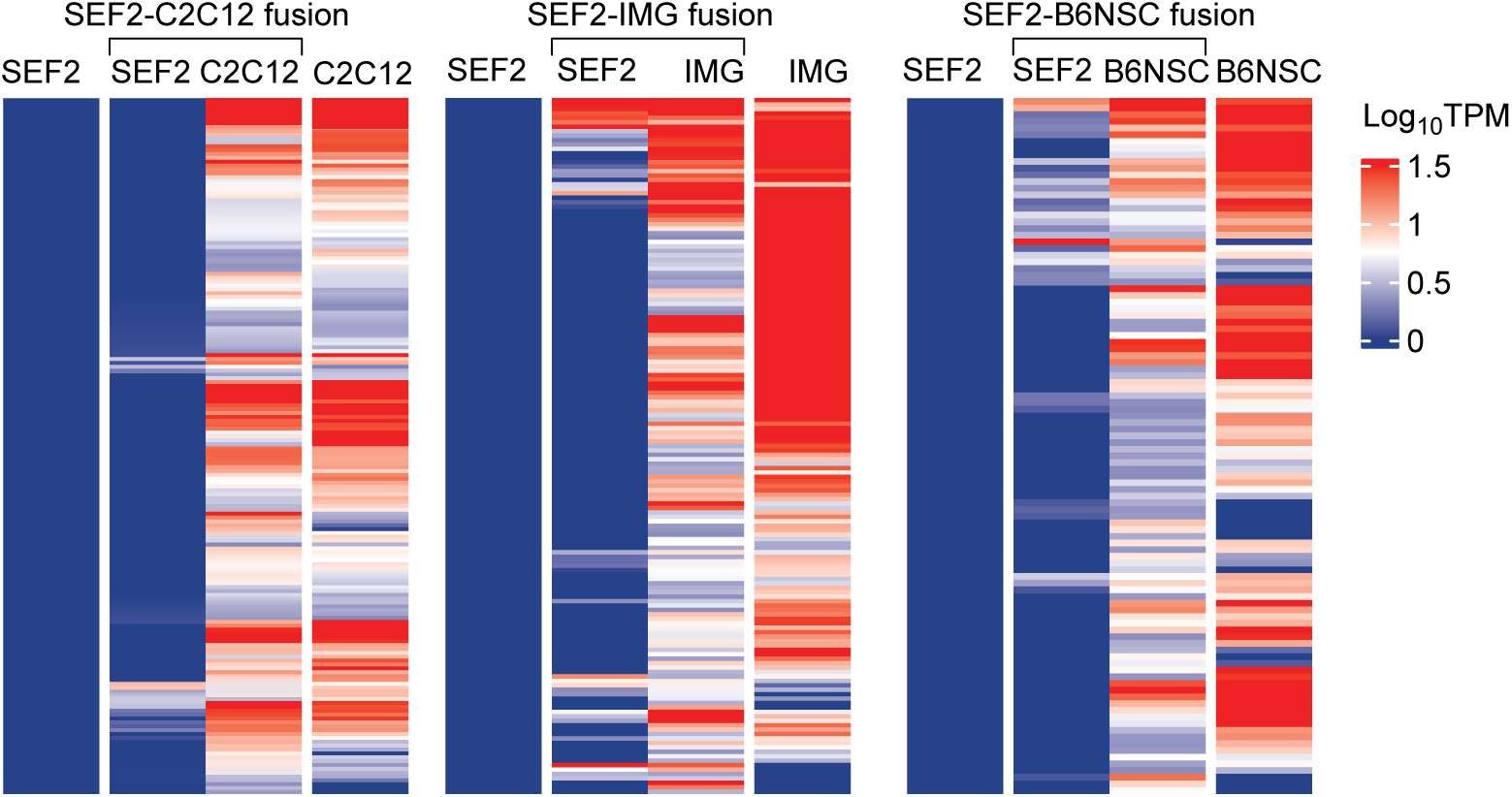
Stability of SEF2 occluded genes in various somatic-somatic fusions. SEF2 genes shown to be occluded in SEF2-C2C12 (n = 180), SEF2-IMG (= 157) and SEF2-B6NSC (n = 104) fusions are combined, with non-extinguished genes in each fusion sample displayed.

**Figure S5.**
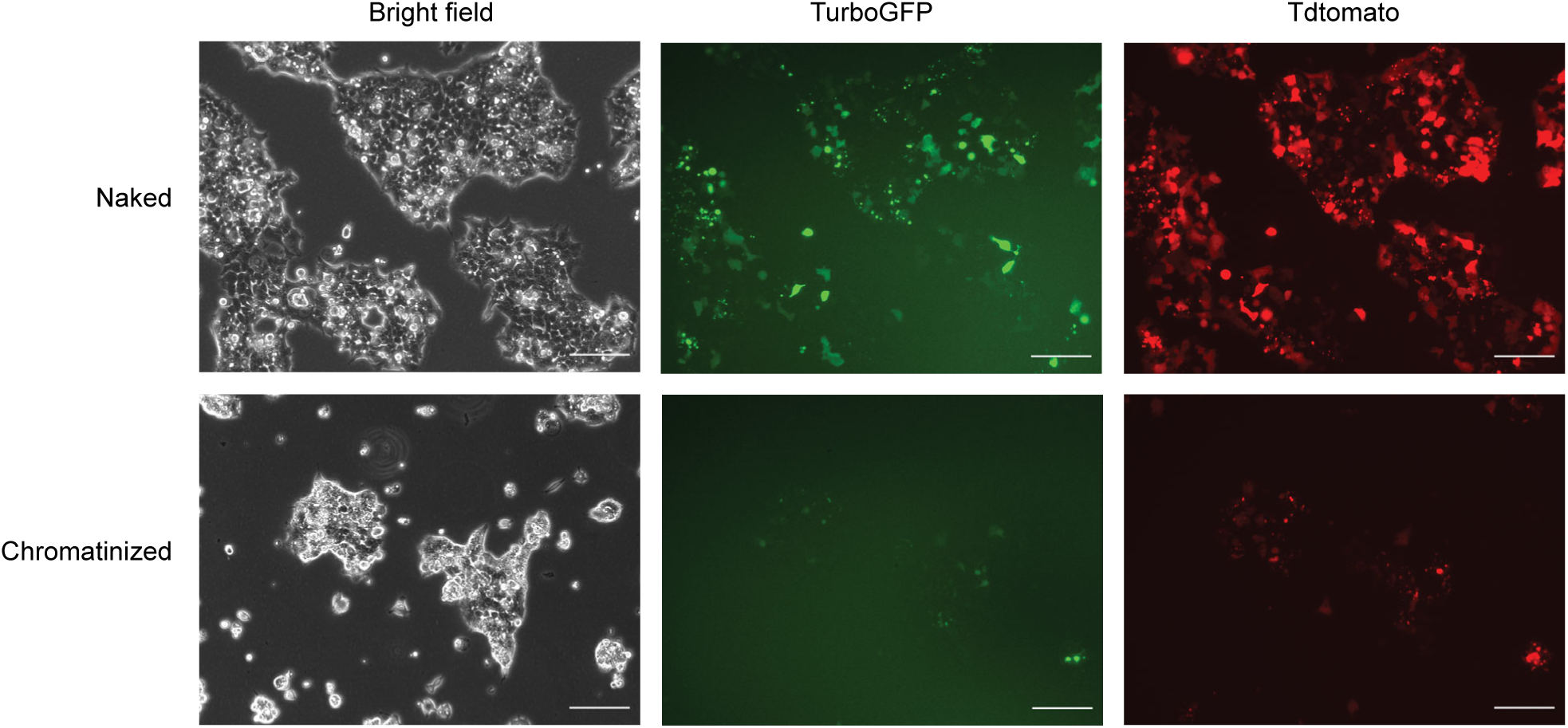
Chromatinized DNA undergoes occlusion. Fluorescence images of 293T cells electorporated with naked or chromatinized plasmids containing Hsp68 driving TurboGFP or EF1A driving Tdtomato. Scale bar: 200 um.

**Figure S6.**
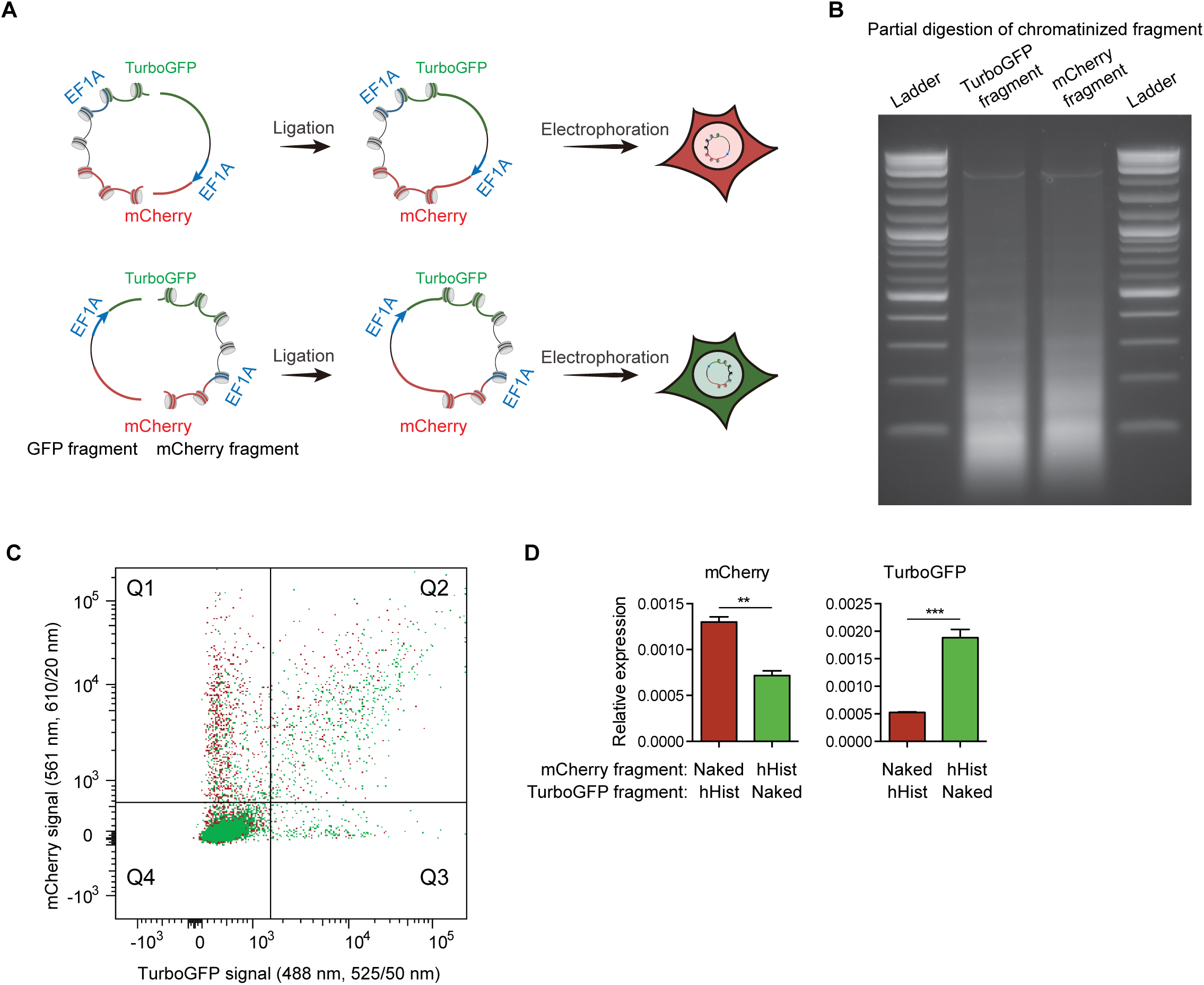
Half-chromatinized episomal DNA is occluded in the chromatinized half. (**A**) Schematic of the half-chromatinization assay. Expression cassettes of TurboGFP and mCherry are split in the middle of the coding sequence. Either the TurboGFP or mCherry fragment is chromatinized while the other one is kept naked. The two fragments are ligated to reconstitute complete TurboGFP and mCherry reporters, followed by electroporation into 293T cells. (**B**) Partial digestion by MNase of half-chromatinized DNA following ligation. (**C**) Flow cytometric analysis of 293T cells electroporated with half-chromatinized DNA. Green dots represent cells containing ligation product of chromatinized mCherry fragment with naked TurboGFP. Red dots represent cells containing ligation product of chromatinized TurboGFP fragment with naked mCherry. (**D**) Relative expression of mCherry and TurboGFP from ligated half-chromatinized DNA. Either the mCherry or TurboGFP fragment was chromatinized with HeLa core histones (hHist), and ligated to the naked form of the TurboGFP or mCherry fragment, respectively. RNA levels were measured by RT-qPCR and normalized to Actb (n = 3, unpaired t-test). Primers were designed across ligation junctions.

**Figure S7.**
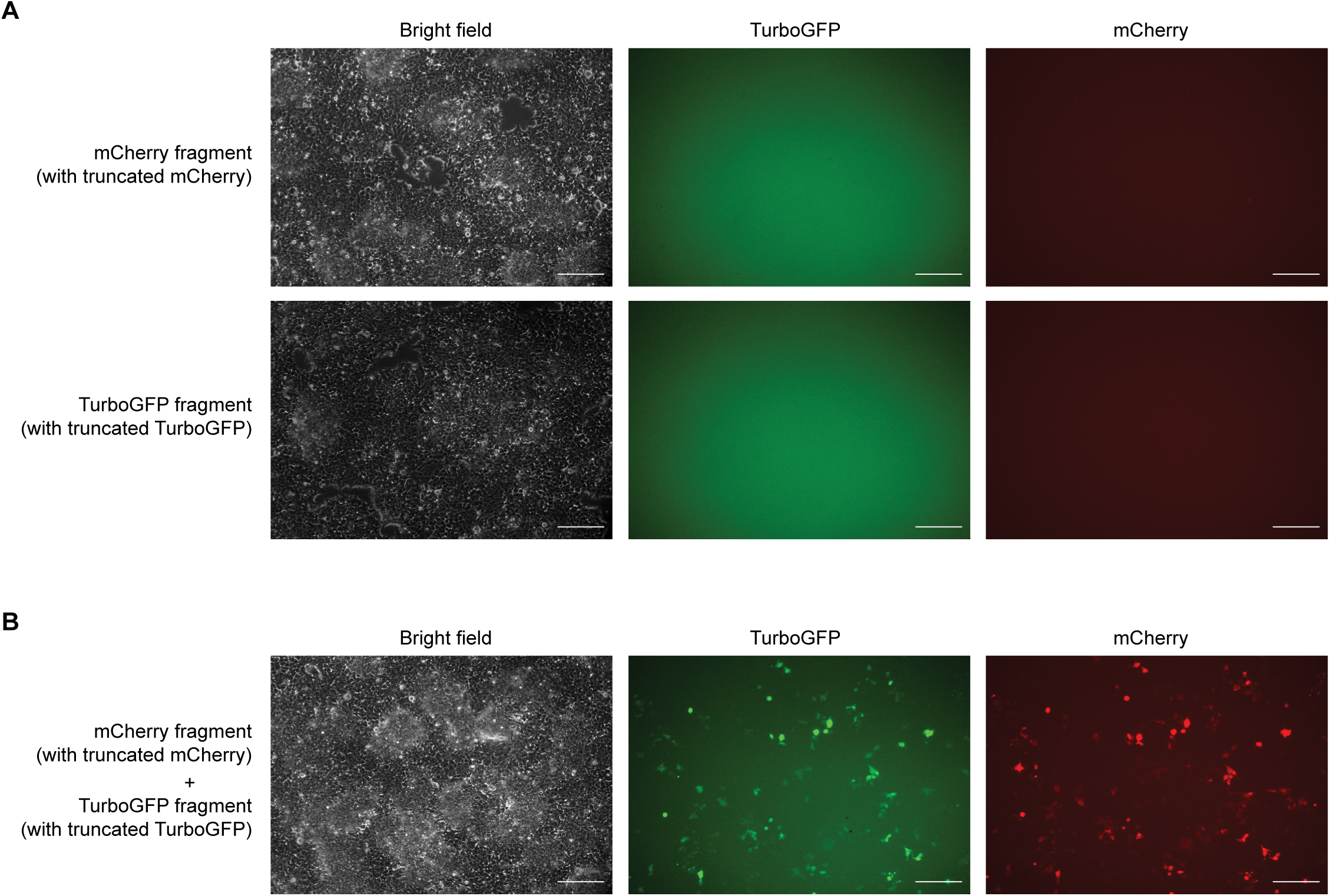
Spontaneous ligation between mCherry and TurboGFP fragments in 293T cells. Expression cassettes of TurboGFP and mCherry were split in the middle of the coding sequence. The two fragments, which had matching sticky ends, were electroporated into 293T cells either individually that failed to produce any fluorescence (A), or simultaneously that produced both TurboGFP and mCherry fluorescence due to spontaneous ligation between the two fragments in cells (B). Scale bar: 200 um.

**Figure S8.**
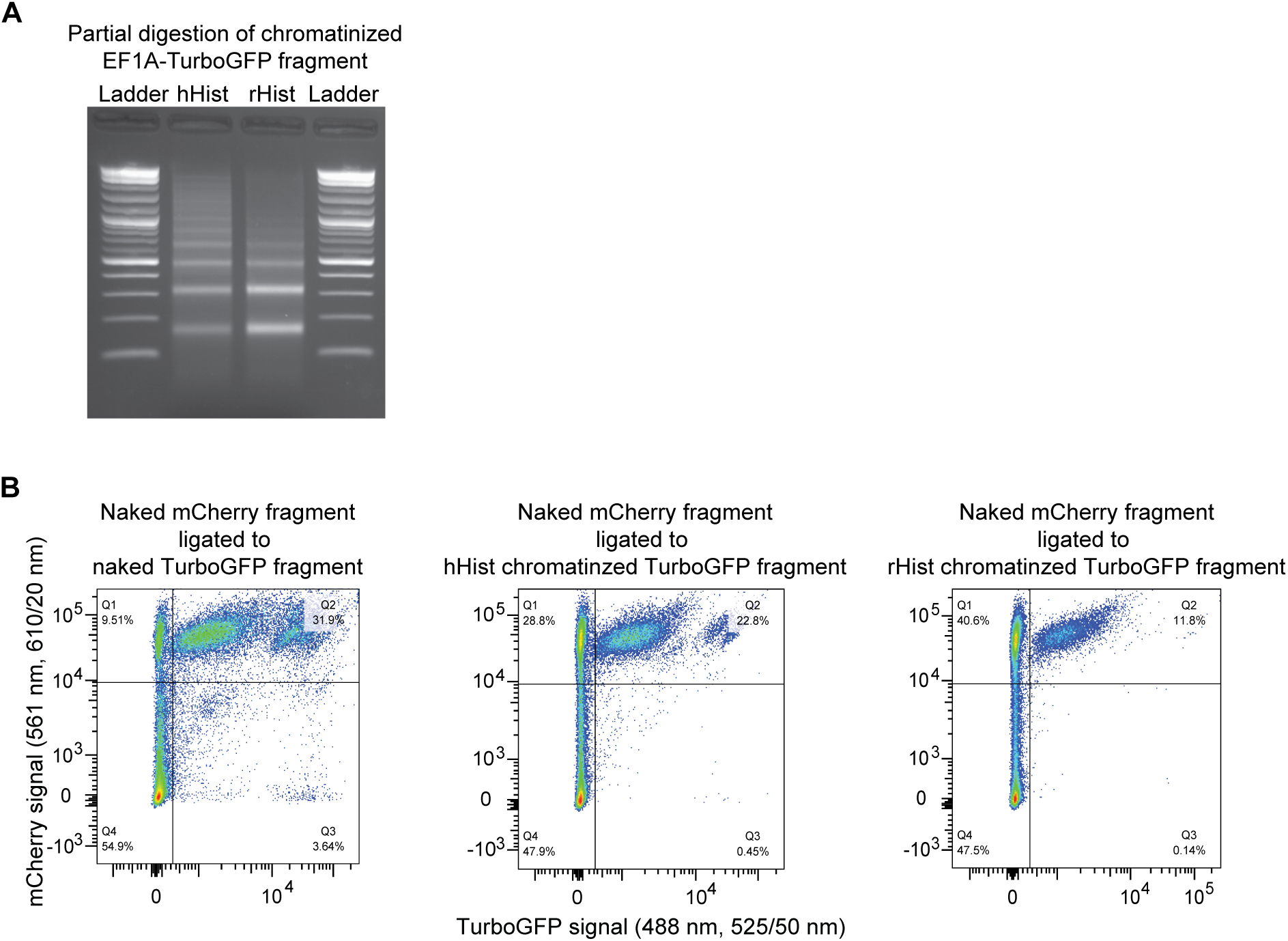
Heritability of nucleosome-mediated gene occlusion through DNA replication. (**A**) Partial digestion by MNase of chromatinized TurboGFP fragment. (**B**) Flow cytometric analysis of 293T cells carrying integrated copies of the naked mCherry fragment ligated with naked, Hela histone (hHist) chromatinzed, or recombinant histone (rHist) chromatinized TurboGFP fragments.

**Figure S9.**
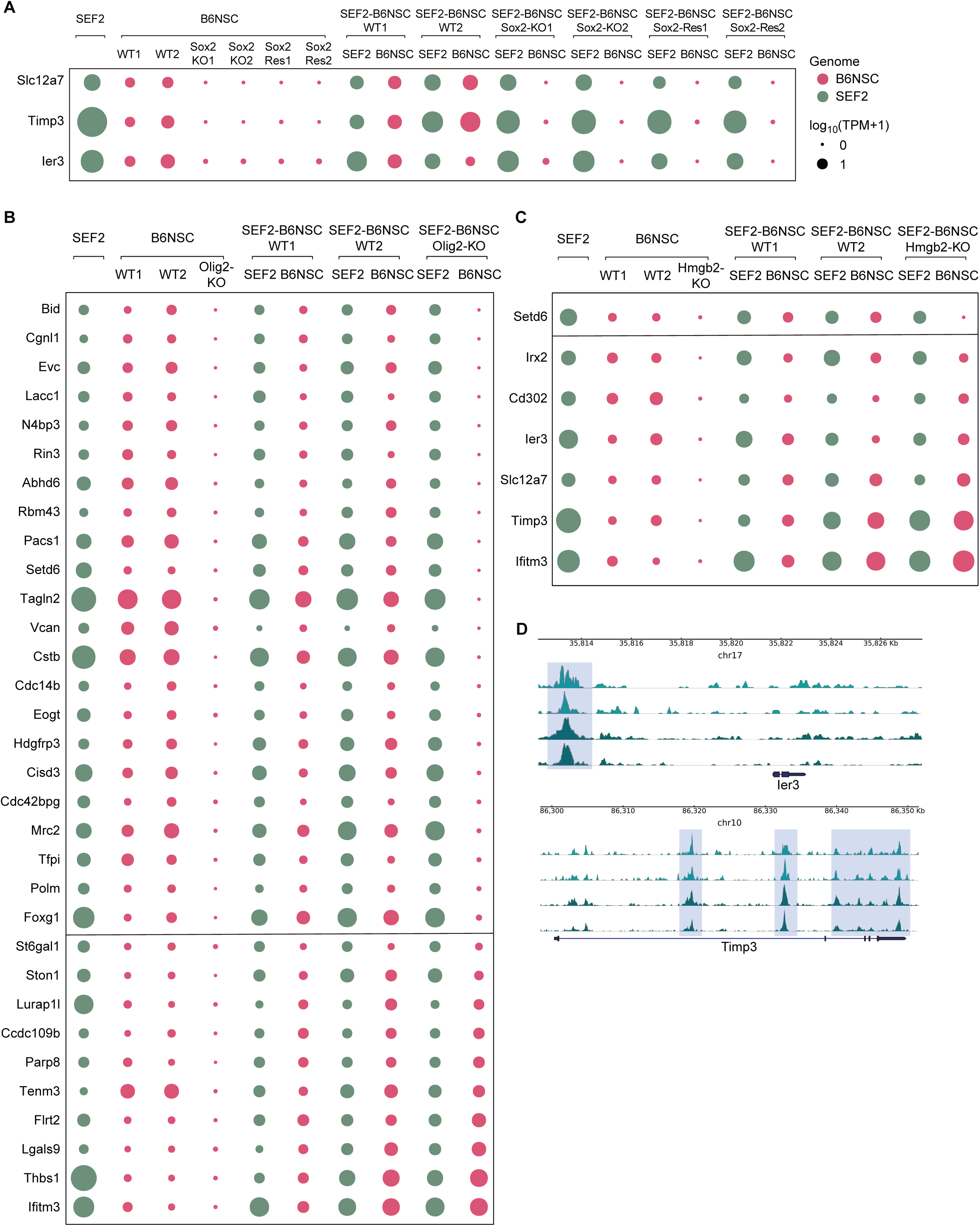
Expressed genes that became silent following Sox2, Olig2 or Hmgb2 knockout. (**A**) Bubble plot depicting expression levels in pre-and post-fusion SEF2 and B6NSC for expressed genes in wildtype B6NSC that became occluded due to Sox2 knockout. Two B6NSC clones homozygous for Sox2 deletion were fused to SEF2 either directly (Sox2-KO1 and Sox2-KO2), or rescued for Sox2 expression by a lentivirus vector expression Sox2 before cell fusion (Sox2-Res1 and Sox2-Res2). Two wildtype B6NSC clones (WT1 and WT2) were used as control. (**B, C**) Bubble plot depicting expression levels for expressed genes in wildtype B6NSC that became silent after Olig2 or Hmgb2 knockout, respectively. (**D**) Sox2 binding profiles of representative expressed genes in B6NSC that became occluded following Sox2 knockout. Sox2 binding was measured by CUT&Tag with a monoclonal antibody (top two tracks for two replicates) and a polyclonal antibody (bottom two tracks for two replicates).

**Figure S10.**
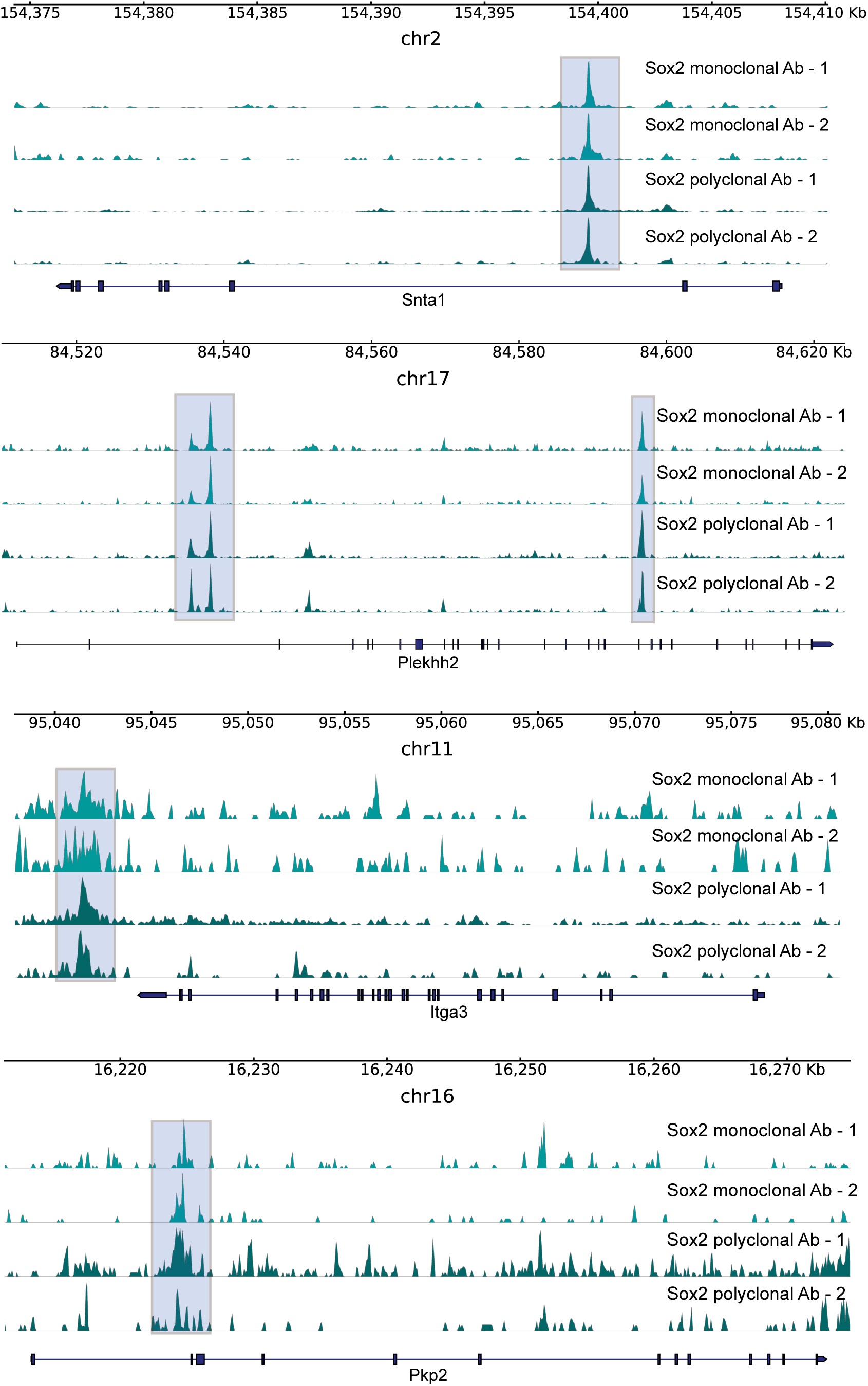
Sox2 binding profiles of activatable genes in B6NSC that became occluded following Sox2 knockout. Sox2 binding was measured by CUT&Tag with a monoclonal antibody (top two tracks for two replicates) and a polyclonal antibody (bottom two tracks for two replicates).

